# Definitive environmental DNA research on aquatic insects: Analysis optimization using the recently developed MtInsects-16S primers set

**DOI:** 10.1101/2023.06.29.547017

**Authors:** Masaki Takenaka, Yuta Hasebe, Koki Yano, Seiya Okamoto, Koji Tojo

## Abstract

Long-term biodiversity monitoring is necessary for the conservation and management of water resources. Notably, aquatic insects have been used as an indicator of water quality because they provide important basic information about freshwater ecosystems and water resources. Although environmental DNA (eDNA) surveys can enable easy and effective biomonitoring of aquatic insects, previous studies have not successfully detected all insect species, and there has been frequent amplification of nontarget taxa (e.g., algae and diatoms). Therefore, we developed a universal primers set, MtInsects-16S, for eDNA analyses of insects in the mtDNA 16SrRNA region. Furthermore, a well-established database of aquatic insects, especially the MtInsects-16S DNA region of Ephemeroptera, Plecoptera, and Trichoptera in Kanagawa Prefecture, which was the target area in this study, was constructed. Therefore, in this study, we conducted eDNA analyses using a universal primers set and using a well-established database. We conducted and compared capture surveys at the same sites to examine the detection capability of eDNA for Insecta. As a result, eDNA analyses using MtInsects- 16S not only detected almost all of the captured species but also detected many more species without amplifying nontarget taxa. This study demonstrated the application of eDNA analyses with unprecedented accuracy and reliability. It was also shown that community structure by eDNA reflected a relatively narrow range at the water sampling point. Although the data accumulation for constructing locally specific databases is an urgent issue, using the MtInsects-16S region is expected to be a breakthrough in the metabarcoding of insects.

## Introduction

Conservation and management of water resources are critical to maintaining a comfortable and prosperous human lifestyle. All the Member States of the United Nations have adopted the 17 Sustainable Development Goals, including Goal 6 (Ensure access to water and sanitation for all) and Goal 15 (Protect, restore and promote sustainable use of terrestrial ecosystems, sustainably manage forests, combat desertification, and halt and reverse land degradation and halt biodiversity loss), which are relevant to inland waters (https://sdgs.un.org/goals). The human impact on freshwater is staggering. More than half of all available freshwater is being used by humans, and water usage continues to increase (Jackson et al., 2001). Thus, freshwater ecosystems are experiencing declines in biodiversity. River ecosystems are particularly critical in terms of biology, which involve various important issues such as adaptive radiation, competition, and speciation (Sekiné & Tojo, 2019; Takenaka & Tojo, 2019; Yano & Tojo, 2020; Almudi et al., 2020).

Notably, aquatic insects have been used as an indicator of water quality because they are sensitive to changes in water quality and environmental health (Cain, et al., 1992; Susmita et al., 2013). Long-term biodiversity monitoring of aquatic insects is important for accumulating basic ecological information for river conservation (Costanza & Mageau, 1999; Sakata et al., 2022). Although collection surveys which involve the capture of aquatic organisms and require a great effort, environmental DNA (eDNA) surveys can enable easy and effective monitoring of the biodiversity of aquatic organisms in many ecosystems (Bohmann et al., 2014; Miya et al., 2015; Deiner et al., 2017; Sakata et al., 2022). Also, eDNA analysis enables the non-invasive assessment of biodiversity and the corresponding genetic diversity for endangered species (Ahn et al., 2020; Sekiya et al., 2017; Yamazaki et al., 2020).

The advancements in metabarcoding primers have greatly accelerated the eDNA studies, especially in fish (Miya et al., 2015). Many previous studies on non-insect groups, particularly those using eDNA, have used primer sets developed from their ribosomal RNA regions (Komai et al., 2019; Miya et al., 2015; Sakata et al., 2022). As for insects, the mtDNA COI region is frequently targeted for DNA barcoding (Folmer et al., 1994). Also, various universal primer sets have been developed to amplify short fragments of the mtDNA COI region for aquatic insects in metabarcoding studies. (Elbrecht et al., 2016; Elbrecht & Leese, 2017; Marquina et al., 2019). For example, Elbrecht and Leese (2017) considered the previously designed primers used to amplify the mtDNA COI region. Although an important advantage of the mtDNA COI region is the availability of an extensive database (Deagle et al., 2014), it is difficult to find a universal primer region for all insects within the mtDNA COI region because no region exists in which the nucleotide sequences match continuously in all insects (Takenaka et al., 2023). On the other hand, the genetic region in the mitochondrial ribosomal RNA region can amplify more species than the mtDNA COI region. This is probably due to a decreased primer bias, especially for algae and bacteria (Elbrecht et al., 2016; Zhang et al., 2020). In insects, eDNA analysis based on the mtDNA COI region often results in the amplification of the non-specific taxa, making it difficult to detect specific eDNA reads (Deiner et al., 2016; Elbrecht & Leese, 2017; Macher et al., 2018; Uchida et al., 2020; Leese et al., 2021). Furthermore, many species cannot be amplified by eDNA analysis based on the mtDNA COI region, and this has not reached a suitable level of accuracy for a diversity of insects (Deiner et al., 2016; Uchida et al., 2020; Leese et al., 2021). Therefore, we developed a universal primers set, MtInsects-16S, for insects in the mtDNA 16S rRNA region, and we confirmed PCR amplification of the target DNA region in all the insect groups that were tested.

One of the reasons why eDNA studies of insects have not been developed further is the lack of databases. We used own database in Kanagawa Prefecture, which is the target area of this study. This database contains the DNA sequences of the mtDNA 16S rRNA regions for many aquatic insects, especially Ephemeroptera, Plecoptera, and Trichoptera. Therefore, we conducted eDNA analyses using this well-established database and established a universal primer set, MtInsects-16S. To investigate the detection ability and accuracy of eDNA analyses using MtInsects-16S, we performed water sampling (and subsequent eDNA analyses) and a capture survey (surveys of insect specimens) at various sites and then compared these results. By eDNA surveys and capture surveys, we confirmed the capability and accuracy of eDNA detection based on MtInsects-16S. Therefore, it is expected to be a breakthrough in the metabarcoding of insects when using the proposed MtInsects-16S set which amplifies these designated genetic regions of the ribosomal mtDNA proposed.

## Materials and Methods

### Study sites

In this study, we conducted actual capture surveys and water sampling at six sites in two river systems (the Sagami-gawa River and Sakawa-gawa River systems) to compare the results of the actual capture survey and eDNA analyses (Fig. 1).

**Figure 1.**
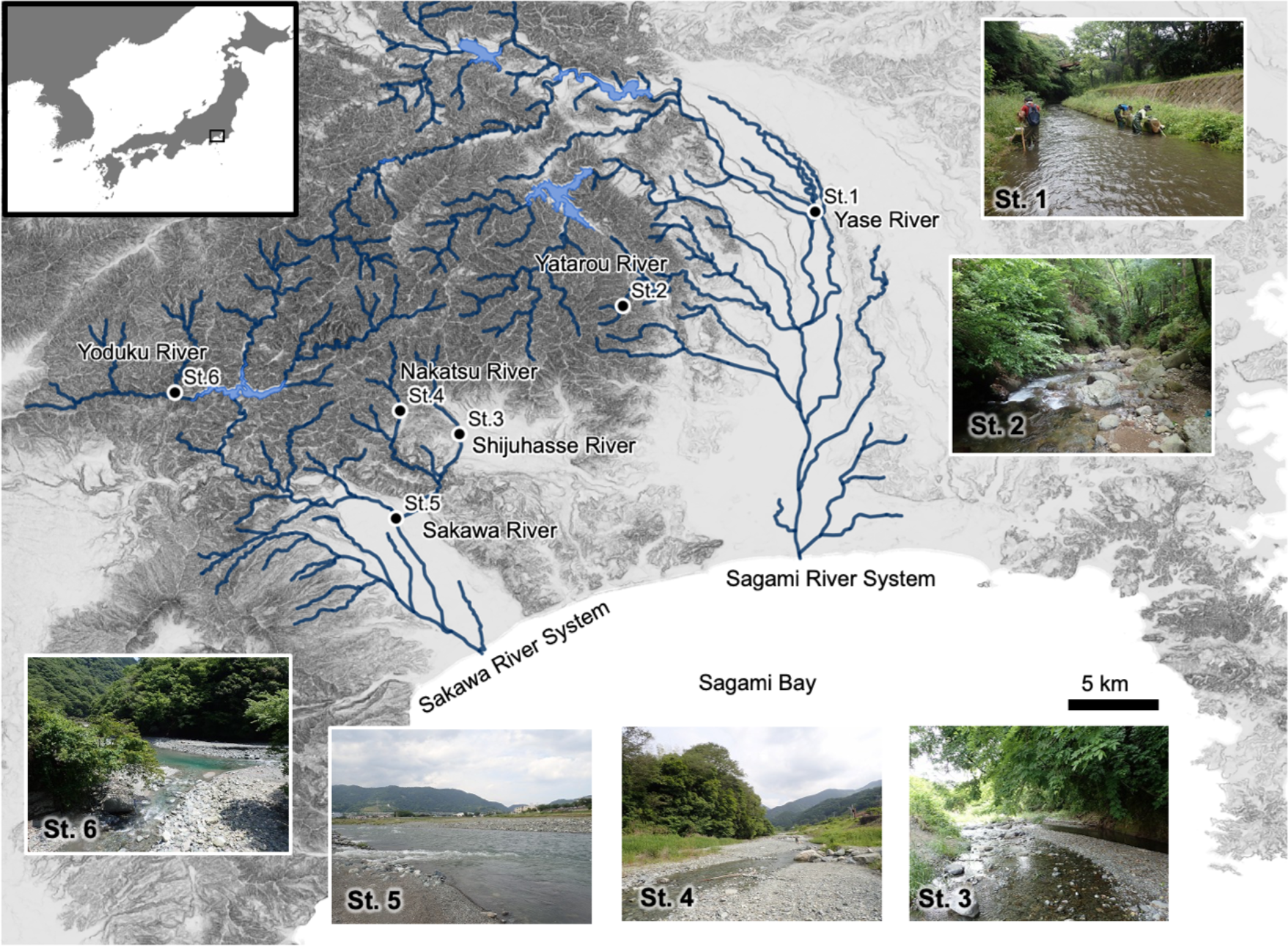
Survey sites. The representative environments of the survey sites are shown at each point.

### Field surveys

Actual capture surveys were conducted on May 24–26, 2022, according to the manual of “National Census on River Environments (NCRE)” reported by the Japanese Government (Administrative Agency of the Ministry of Land, Infrastructure, Transport and Tourism). Regarding the quantitative sampling at each of the six sites, three quadrats (0.25 m x 0.25 m) were set up in lotic water (riffle) environments, using a surver net (net width of 250 mm, and mesh size of 0.493 mm: NGG38), after which qualitative sampling was also conducted using a D-frame net to cover almost all the microhabitats.

### Environmental DNA sampling and sequencing

Filtration and eDNA extraction were performed according to the Environmental DNA Sampling and Experiment Manual ver. 2.1 (The eDNA Society, 2019). At each site, 1 L of surface water was collected for eDNA analyses using a new polypropylene bottle, before the capture surveys were conducted. After water sampling, BAC (final concentration = 0.1%) was added to prevent DNA degradation (Yamanaka et al., 2017), and the samples were stored in a cooler box and filtered within 48 hours.

Each water sample was filtered through Sterivex (0.45 μm pore size, Merck & Co., Inc., Massachusetts, USA) and 2 mL of RNAlater™ Stabilization Solution (Thermo Fisher Scientific, Massachusetts, USA) was added for preservation. Then, the filters were stored at −20℃. During filtration, disposable instruments were used for all parts that directly touched the sample to prevent contamination. Also, 1 L of distilled water was filtered as the negative control and used as a blank sample. Extraction of eDNA from the Sterivex filters was performed according to Wong et al. (2020).

RNAlater was removed using 50 mL syringes (SS-50LZ, Terumo). A lysis buffer mix (PBS 990 μL, Buffer AL 910 μL, Proteinase K 100 μL) was injected and incubated at 56℃ for 30 minutes. Then, the lysis buffer mix was collected using 50 mL syringes (SS-50LZ, Terumo), and 1 mL of molecular biology grade ethanol (99.5%, Fujifilm Wako Pure Chemical Corporation, Osaka, Japan) was added to the lysis buffer mix. The mixture was then purified using a DNeasy Blood and Tissue kit. Finally, the column was eluted with 150 μL of AE buffer.

Each total genomic eDNA sample was used to library prep for high-throughput sequencing using a MiSeq platform (Illumina, San Diego, CA, USA). The primers set, MtInsects-16S, was used to amplify DNA fragments [the mitochondrial DNA (mtDNA) 16S rRNA region]. MtInsects-16S (F: 5’ – GGA CGA GAA GAC CCT WTA GA – 3’ and R: 5’ – ATC CAA CAT CGA GGT CGC AA – 3’) is a primers set for DNA barcoding that was recently designed in a previous study (Takenaka et al., 2023). Library preparation was conducted in two patterns, 2-step PCR and 3-step PCR. In general, library preparation for high-throughput sequencing is conducted through 2-step PCR. However, when conducting PCR on DNA extraction samples using primers with adapter sequences, amplification bias is known to occur, which is specific to the template sequence (O’Donnell et al., 2016; Mansfeldt et al., 2020). In this case, it is performed a preparatory PCR (0th PCR) using primers with no adapter sequences (only primers sequences) before the 1st PCR using primers with adapter sequences (O’Donnell et al., 2016).

Library preparation by 2-step PCR was performed as follows. The 1st PCR was conducted using the KOD One PCR Master Mix (TOYOBO, Osaka, Japan) with primers and adapter sequences (F: 5’ – ACA CTC TTT CCC TAC ACG ACG CTC TTC CGA TCT NNN – 3’ and R: 5’ – GTG ACT GGAGTT CAG ACG TGT GCT CTT CCG ATC TNN N, all adapter sequences in this study are the same) to assign the adapter region for the 2nd PCR. Four replicates of each sample were run to minimize PCR dropouts. The following PCR protocol was used: 98℃ for 10 sec; 35× (98℃ for 10 sec, 55℃ for 5 sec, 68℃ for 5 sec). A blank sample prepared during DNA extraction was used as a negative control, and PCR was performed using the same protocol. Four replicates of PCR products were pooled and 40 μL was subjected to 0.6×–0.8× Right Side Size Selection by SPRI select for purification and removal of nontargeted amplification. Then, the DNA concentration was confirmed using the 4150 TapeStation system (Agilent) and diluted to 100 pg/µL. The 2nd PCR was also conducted using the KOD One PCR Master Mix (TOYOBO, Osaka, Japan) for assigning sequences an index to identify individual samples and binding to the flow cell of the high-throughput sequencing sequencer. The following PCR protocol was used: 10× (98℃ for 10 sec, 60℃ for 5 sec, 68℃ for 5 sec). PCR products were subjected to 0.6×–0.8× Right Side Size Selection by SPRI select for purification and removal of nontargeted amplification.

Library preparation by 3-step PCR was performed as follows. First, the 0th PCR was performed using the KOD One PCR Master Mix (TOYOBO, Osaka, Japan) with only primer sequences (no adapter sequences). Four replicates of each sample were run to minimize PCR dropouts. The following PCR protocol was used: 98℃ for 10 sec; 26× (98℃ for 10 sec, 55℃ for 5 sec, 68℃ for 5 sec). A blank sample prepared during DNA extraction was used as a negative control, and PCR was performed using the same protocol. Four replicates of PCR products were pooled. The 1st PCR and the 2nd PCR were conducted using the same protocol, but only 10 cycles were performed for the 1st PCR.

Sequencing of the DNA library was outsourced to Bioengineering Lab. Co., Ltd. A sequence was performed using the MiSeq platform (Illumina, San Diego, CA, USA) with approximately 20% of the Reagent Kit v3 (2x300 bp). All raw sequencing reads were registered with the DNA Data Bank of Japan (DDBJ: PRJDB15936).

### Bioinformatics

For preprocessing the raw sequencing reads, we used Usearch v11.0.667(https://drive5.com/usearch/download.html) and cutprimers.py in Python (https://github.com/aakechin/cutPrimers). Paired-end reads were merged from the MiSeq platform raw sequencing reads using the default settings. The sequences after bases with a Phred Quality Score of 2 and paired reads with too many mismatches in the aligned region (> 5 bp or less than 90% identity) were discarded. Then, read quality filtering was performed using Usearch’s “fastq_flter” command with a sum of prediction errors threshold of < 1.0 for all bases in the read and read lengths of >180 bp. After that, cutPrimers.py was used to remove the primer sequences, and the number of preprocessed reads was calculated for each unique base sequence using the “fastx_uniques” command. Finally, using Userarch’s “unoise3” command, we checked for PCR errors and chimeric reads and performed error correction for discarded amplicon reads, according to the procedure reported by Edgar and Flyvbjerg (2015).

Blank processing was performed for ZOTU (zero-radius OTU). Specifically, we subtracted the number of blank ZOTU reads prepared for each library preparation of 2-step PCR and 3-step PCR from the number of ZOTU reads for each sample. At the species level, there is a possibility that the number of detected reads can affect the rate of agreement with the actual capture survey, so the number of reads at each survey site was adjusted to match the site with the lowest number of reads. Then, ZOTUs with 7 or fewer reads were treated as no data.

A BLAST search was performed on the final reads using two databases: local BLAST 2.13.0+ (https://ftp.ncbi.nlm.nih.gov/blast/executables/blast+/LATEST/) and the DNA database for insects created by Kanagawa Prefecture (https://www.pref.kanagawa.jp/docs/b4f/suigen/edna.html), which was constructed mainly by one of the authors (YH) for eDNA analyses of insects, and the GenBank (https://blast.ncbi.nlm.nih.gov/Blast.cgi). We employed three thresholds for taxonomic identification.

The first threshold was the e-value of 1e^-40^ for the phylum level. The e-value of 1e^-20^ separated three groups: Insecta *s. str.*, Diplura, and Collembola. Therefore, we used the e-value of 1e^-40^, considering a certain degree of certainty. Within Insecta *s. str.*, the second threshold was the e-value of 1e^-60^ for the order level (p-iden > 90%, q-cover > 90%). In this identification, if the prefectural DB and GenBank were judged to be different, the data were treated as invalid data.

The third threshold was set as p-ident×q-cover ≥ 0.98 for the species level. This analysis is the most important because it is necessary to strictly identify the species. If the prefectural DB and GenBank were judged to be different, the higher number above the threshold was used, but when the numbers were the same, the species were listed together. Genus-level sequences, such as ’sp.’ in the database, were treated as invalid data. In this analysis, we targeted aquatic insects of Ephemeroptera, Plecoptera, and Trichoptera for comparison with the actual capture surveys because these orders are major organisms in the rivers and because the prefectural database almost covers the DNA sequences of the species confirmed at the survey sites. Specimens that could not be identified by nymph/larval morphological characteristics in the actual capture surveys were treated as one taxonomic group, and the eDNA reads were also treated as single species. Similarly, species that could not be distinguished by eDNA were also treated as one taxonomic group (“*Epeorus latifolium*/*Epeorus l-nigrum*/*Epeorus napaeus*,” “*Drunella sachalinensis*/*Drunella kohnoi*,” “*Ephemera strigata*/*Ephemera japonica*,” “*Nemoura fulva*/*Nemoura redimiculum*,” “*Sweltsa nikkoensis*/*Sweltsa kibunensis*,” “*Hydropsyche selysi*/*Hydropsyche gifuana*,” “*Rhyacophila towadensis*/*Rhyacophila lezeyi*,” “*Rhyacophila nipponica*/*Rhyacophila nigrocephala*”). To exclude accidental data due to flow down, quantitative surveys with only 1 sample and/or qualitative surveys with less than 5 samples were treated as non-data.

### PCR inhibition by adapter sequences

To determine the genomes of aquatic insects for adapter sequences that can be used in high-throughput sequencing, we downloaded genome data sets for ten species from Ephemeroptera, Plecoptera, and Trichoptera from GenBank. Using the obtained genomic information, we searched for the presence of DNA regions that were similar to the adapter sequences using local BLAST 2.13.0+ (https://ftp.ncbi.nlm.nih.gov/blast/executables/blast+/LATEST/).

### Environmental factors measured by GIS (Geographic Information System)

To evaluate the environmental factors of the six sampling sites, the following factors were measured using a Geographic Information System (GIS). (1) Altitude: the underlying elevation data were obtained from a 5 m digital elevation model (DEM) provided by the Geospatial Information Authority of Japan. (2) Strahler stream order: the stream order (Strahler, 1957) data were calculated based on a 1/25,000 scale map obtained from the National Land Numerical Information, provided by the Geospatial Information Authority of Japan. (3) Riverbed slope degree: the degree of riverbed slope was calculated by GIS mapping based on the distance between points at which there was a 10 m elevation difference, with the sampling site set as the base point, using the 5-m DEM provided by the Geospatial Information Authority of Japan.

### Community structure analysis

To compare the aquatic insect community structures among the survey sites, hierarchical cluster analysis and non-metric multidimensional scaling (nMDS) were performed based on Jaccard distances (presence/absence data) (hierarchical cluster analysis, *hcluster* function in R package *cluster*; nMDS, *metaMDS* function in R package *vegan*) (Oksanen et al., 2020). The software R ver. 4.2.1 was used for this analysis (R Core Team, 2022).

## Results

Within all the reads detected by eDNA analyses, the sequences were used to create OTUs for the e-value of 1e^-40^. As a result, 4,843,866 bp reads were clustered into groups, and 113,579 bp reads were not aligned (Table S1: 547,479–973,073 bp/survey site). Then, we examined the percentage of taxa detected within all the reads obtained. At each site, reads for Insecta *s. str.* accounted for more than 90%, except for St. 2 (Table S1; Fig. 2). For St. 2, relatively more reads of Hydrozoa and Osteichthyes were detected, in addition to Insecta *s. str.* (Table S1). Moreover, less than 0.1% of the reads detected at the sites were algal or bacterial (Table S1).

**Figure 2.**
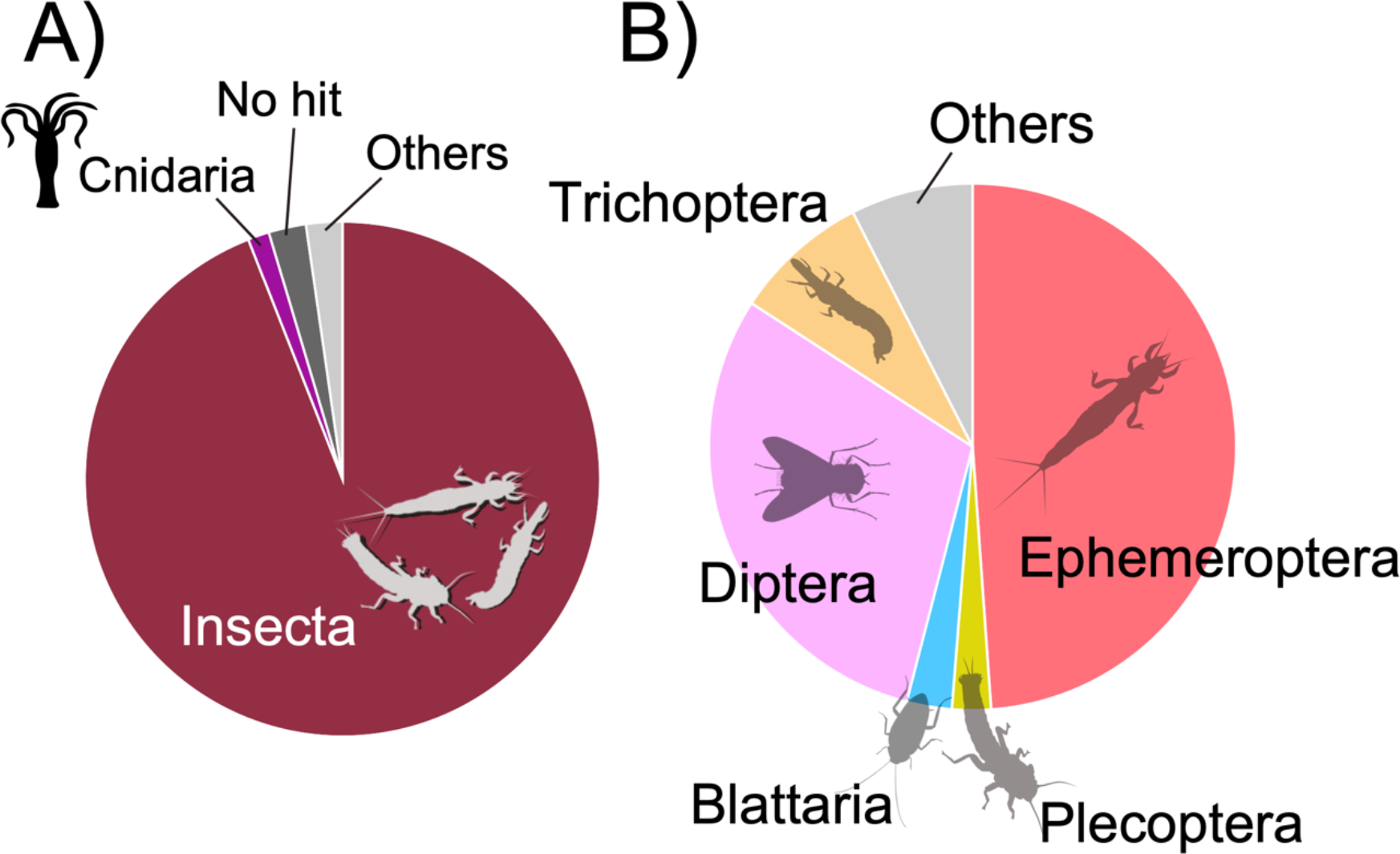
Pie charts showing the percentage of reads detected from eDNA analyses in summarize of all survey sites. A) Composition ratio of major animal taxa (No hit, eDNA reads that did not hit the database; Others, detecting eDNA read of other groups), B) Composition ratio at order level within Insecta.

A similar analysis was performed at the order level within the class Insecta. This analysis was used to create OTUs for under the e-value of 1e^-60^. As a result, 4,230,728 bp reads were clustered into groups (Table S2: 516,629–864,418 bp sequence site), and most of the reads detected by eDNA analyses were from the orders Ephemeroptera, Diptera, and Trichoptera (Table S2; Fig. 2).

### Comparing actual capture survey and eDNA analyses

To examine the accuracy and detection capability of eDNA, we compared the results of actual capture surveys for benthic invertebrates with the eDNA analyses of water samples from the six sites. The detected species are listed in Table S3. Analyses of eDNA were detected 44 species from 8 families of Ephemeroptera, 31 species from 6 families of Plecoptera, and 42 species from 16 families of Trichoptera. From the actual capture surveys, 38 species from 8 families of Ephemeroptera, 6 species from 2 families of Plecoptera, and 23 species from 11 families of Trichoptera were collected.

For each survey site, the collected species and the numbers of individuals collected in three quantitative and qualitative surveys are shown in Table S3, and the number of reads for each species detected by eDNA analyses is also shown. In this analysis, 2-step and 3-step PCR (i.e., double and/or triple PCR) were conducted (Table S3). As a result of the eDNA analyses by 2-step PCR, 42.3% of the species matched the capture surveys. The eDNA analyses detected 96.2% of the total species. Thus, only 3.8% of the species were undetected by eDNA analyses, detected by only actual capture surveys. The detection rate of eDNA slightly increased for all sites analyzed by 3-step PCR (Table S3).

Regarding the separate analyses performed for Ephemeroptera, Plecoptera, and Trichoptera, only a few species were not detected by eDNA analysis (Table S3). Of the three insect orders, only 8 species were collected by actual capture surveys but undetected by eDNA analysis. Among them, 6 species (*Baetis* sp. J, *Baetis taiwanensis*, *Labiobaetis atrebatinus orientalis*, *Ecdyonurus kibunensis*, *Rhithrogena tetrapunctigera*, *Drunella ishiyamana*) were detected from other survey sites by eDNA analysis. However, for the remaining two species (*Uenoa tokunagai*, *Gumaga orientalis*), no eDNA reads were detected from any of the sites.

### PCR inhibition by adapter sequences

After searching for sequences similar to the adapter sequences on the genome of various aquatic insects, some regions that have high similarity and low coverage compared with the adapter sequences were detected in almost all the species genomes (approximately 50% of query cover and 100% of per- identification values: Tables S4, S5). There were no similar sequences found for *Limnephilus maemoratus*. Moreover, we searched for homologs of all the adapter-like regions by performing a BLAST search using the GenBank database (https://blast.ncbi.nlm.nih.gov/Blast.cgi); however, 16S rRNA was not observed in any of the top hits.

### Community structure assessment of aquatic insects using eDNA

The community structure at each site was compared using species presence/absence data as detected by eDNA analyses via 3-step PCR. As a result, the sites were divided into two clusters based on their similar community structures (St. 1 and 5; and St. 2, 3, 4, and 6), which were represented by different environments (Fig. 4). Although St.1 and 5 exhibited a low riverbed slope gradient and a low altitude (relatively downstream characteristics), St.1 was classified as an upstream environment owing to its stream order of 1. The other cluster (St. 2, 3, 4, and 6) had a high altitude, steep riverbed slope degree, and large Strahler stream order (Fig. 4).

**Figure 3.**
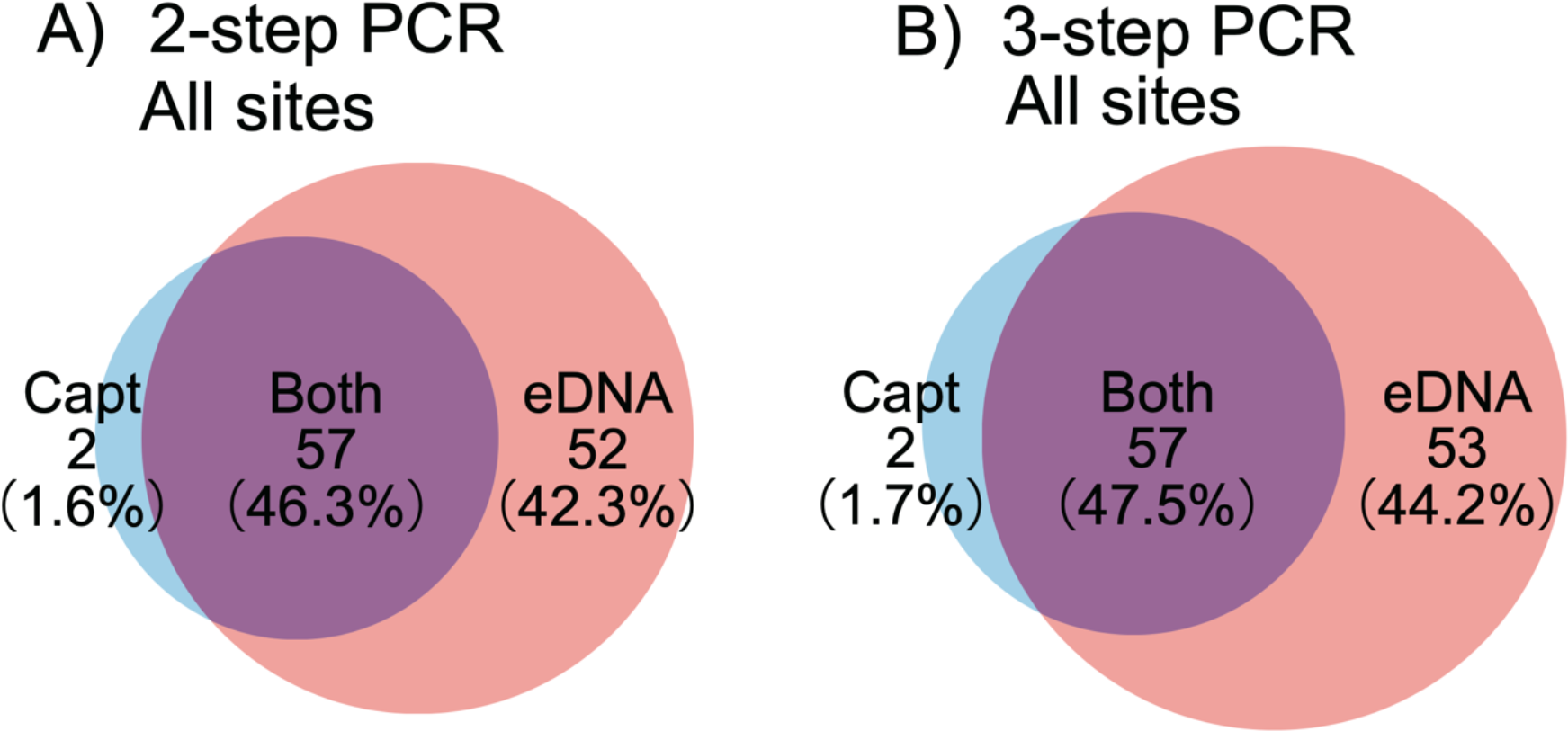
Venn diagrams for comparison of the number of species detected by eDNA analyses and actual capture surveys, and comparison of two library prep patterns. A) 2-step PCR and B) 3-step PCR.

**Figure 4.**
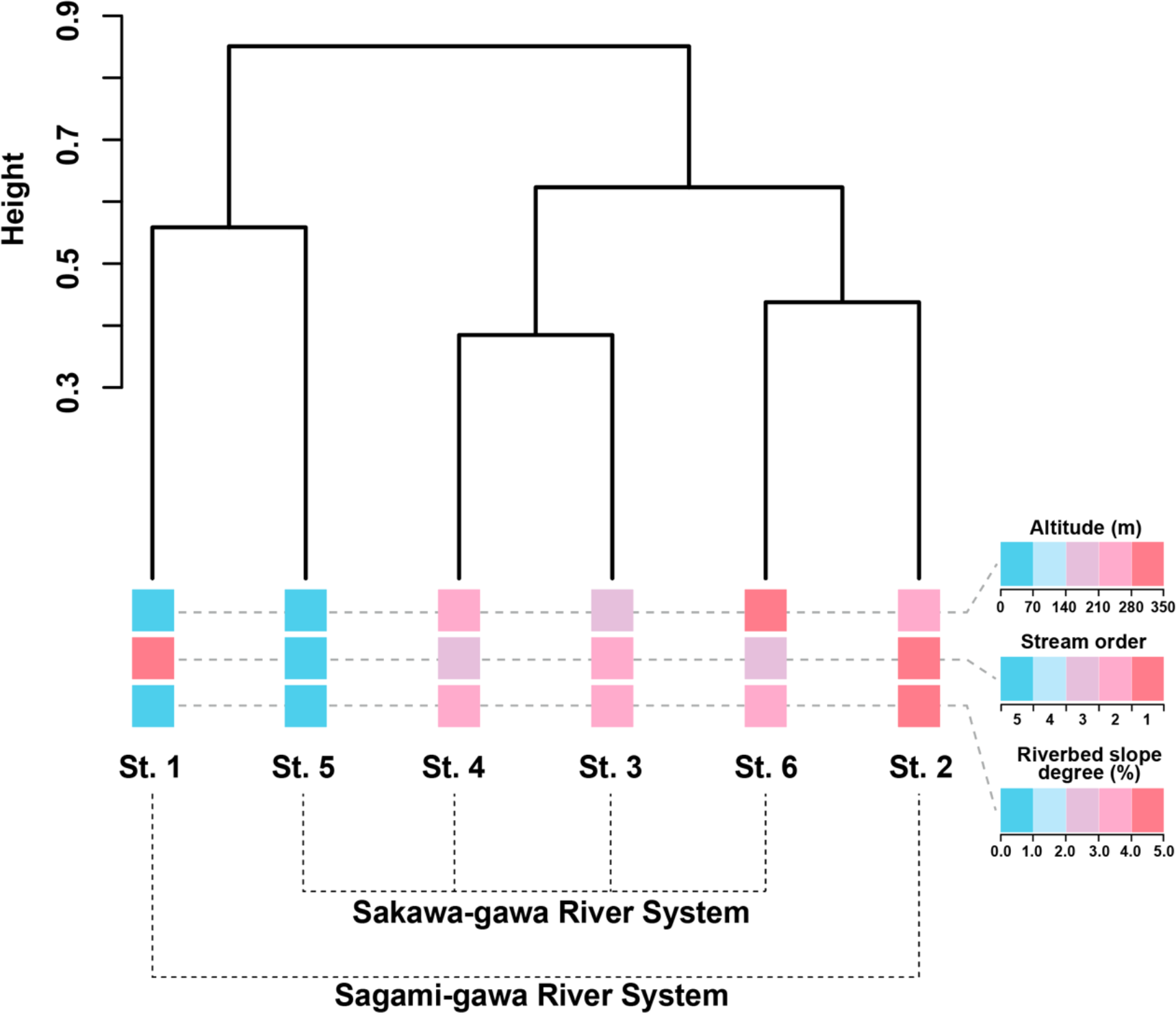
Community structures of aquatic insects detected by eDNA analyses. Hierarchical clustering of two river systems (Sagami-gawa and Sakawa-gawa River systems) and the similarity of their community structures. Cluster dendrogram shows the community structure of aquatic insects. The environments of each site are shown under the name of each site on the dendrogram using a five-grade evaluation of the altitude, stream order, and riverbed slope.

## Discussion

The eDNA analysis method of aquatic insects is not only a breakthrough tool for the assessment of river biodiversity and genetic diversity but also the assessment of river water quality. For fish and other organisms, the design of suitable universal primers for metabarcoding analysis has enabled great progress in the development of eDNA analysis methods. However, the eDNA analysis of insects remains underdeveloped. In particular, insects that are known to be the richest in species diversity on earth (Grimaldi & Engel, 2005; Tojo et al., 2017) have generally been difficult to analyze using a uniform method across all insect species. In our recent study, we designed a suitable primers set for DNA barcoding, MtInsects-16S, which can amplify DNA for all major clades of insects (Takenaka et al., 2023).

### High detection capability of eDNA for Insecta

Metabarcoding analysis has typically been performed in the mtDNA COI region in insects, but eDNA analysis using the mtDNA COI region has not provided sufficiently high detection (Deiner et al., 2016; Macher et al., 2018; Uchida et al., 2020; Leese et al., 2021). The advantage of the mtDNA COI region is the high availability of referenceable data from multiple databases such as GenBank and BOLD system (Leese et al., 2021; Takenaka et al., 2023).

Although the LCO1490 and HCO2198 primers set (Folmer et al., 1994) has often been used in insects and for eDNA analysis based on the mtDNA COI region, few insect eDNA reads have been detected (Deiner et al., 2016; Uchida et al., 2020). Uchida et al. (2020) reported that a low rate of insect eDNA reads was detected by eDNA analyses. It is also reported that, the eDNA detection rate of insects has been poor in studies using the BF2 and BR2 primers set (Elbrecht & Leese, 2017), which are also used often in studies of insects based on the mtDNA COI region. Leese et al. (2021) reported that Arthropoda, including Insecta, accounted for only about 24% of the reads detected by eDNA analyses using the BF2 and BR2 primer set, and Macher et al., (2018) reported that Metazoa, including Insecta, accounted for only about 20% of the reads detected by eDNA analyses. Similarly, only 12% of the eDNA reads were found among benthic invertebrate taxa (Gleason et al., 2020). Such an efficient analysis for abundant nontarget taxa, such as algae, bacteria, and fungi, has not been reported. This is partly because the penalty scores for abundant nontarget taxa, such as algae, bacteria, and fungi, were substantially higher (Leese et al., 2021).

Therefore, Leese et al., (2021) designed a new set of PCR primers (fwhF2 and EPTDr2n) that avoids problems related to the amplification of nontarget taxa and reported that a high level of detection was achieved for aquatic insects. More species were detected using eDNA analyses of water sampling in rivers using the new set of primers than all the species previously collected over 20 years of long-term monitoring (Leese et al., 2021). However, using the fwhF2 and EPTDr2n primers set, nearly half of the species collected during long-term monitoring were not detected by eDNA analyses (Leese et al., 2021). Many species have likely been missed by eDNA surveys; however, this remains uncertain because a comparative actual capture survey has not been conducted during the same period. Also, there are differences in seasonal patterns over 20 years, and misidentification is also possible.

In this study, we conducted eDNA analyses using a new primers set, MtInsects-16S (Takenaka et al., 2023). As a result, approximately 90% of the eDNA reads detected in the eDNA analyses were from the Insecta class. The next most detected eDNA reads were from Cnidaria. Cnidaria eDNA reads have been the most easily detected using the BF2 and BR2 primers set (Elbrecht & Leese, 2017), which is widely used in the mtDNA COI region (Elbrecht et al., 2016). This suggests that Cnidaria account for a relatively large proportion of biomass existing in rivers.

PCR is generally performed twice for metabarcoding analysis (i.e., 1st tailed PCR and then 2nd tailed PCR), before sequencing. To attach an index for analysis by high-throughput sequencing, the adapter sequence is attached to the primer for the 2nd PCR during the 1st PCR. However, regions similar to the adapter sequence often exist within the genomes of various Ephemeroptera, Plecoptera, and Trichoptera species, and amplification bias can occur when conducting PCR on DNA extraction samples using primers with adapter sequences (O’Donnell et al., 2016; Mansfeldt et al., 2020). Therefore, regions that are similar to the adapter sequence can inhibit the PCR. In this study, DNA regions were found that resemble the adapter sequences on the genomes of each species in Ephemeroptera, Plecoptera, and Trichoptera; however, we did not confirm whether they affected the results. Instead, we conducted a pattern in which the 0th PCR is performed before the 1st PCR. The results of this 3-step PCR procedure were compared with those of the standard 2-step PCR (i.e., 1st PCR and 2nd PCR). As a result, the detection rate of eDNA by 3-step PCR was higher than that obtained by 2-step PCR, for all analyses (Fig. S3), including the detection rate of eDNA in Ephemeroptera, Plecoptera, and Trichoptera (Fig. S4). Still, it is not known whether regions similar to the adapter sequences affected the PCR. It is necessary to consider not only the detection of these species but also how the number of detected reads is affected for each species. In particular, the amplification biases should be addressed when considering the detection rate of eDNA in quantitative analyses.

### Highly accurate eDNA analyses of aquatic insects

Regarding the BF2 and BR2 primers set commonly used for metabarcoding in mtDNA COI region (Elbrecht & Leese, 2017), the fwhF2 and EPTDr2n primers set developed to avoid amplification of nontarget taxa (Leese et al., 2021), and the LCO1490 and HCO2198 primers set used for DNA in general (Folmer et al., 1994), a consensus has not been established as to which set or method is the most effective. In this study, the sequences were used to create OTUs based on 98% sequence identity and 98% query cover within Insecta. As a result, eDNA analyses not only detected almost all of the collected species but also detected more species, other than those that were collected. An outcome has never been reported in which almost all of the collected species were detected by eDNA analyses of water samples. Only 8 collected species were undetected by eDNA at some study sites, but 6 of those species were detected at other sites. Therefore, it is likely that eDNA was not contained in the sampled water of such sites, especially considering the small biomass in the river, rather than being the result of unsuccessful PCR amplification.

However, 2 species that were collected in the quantitative survey were not detected by eDNA analysis. These species may not be detectable using the MtInsects-16S primers set. Notably, *Uenoa tokunagai* (wtr-14_CMG) and *Gumaga orientalis* (OL678018, NC_069258) had two-base pair mismatches with MtInsects-16S_F but matched perfectly with MtInsects-16S_R. Therefore, it is also possible that eDNA was not contained in the sampling water because of the small biomass in the river. In the future, it is necessary to investigate these undetected species in more detail.

### Community structure of aquatic insect by eDNA

Finally, we compared community structures based on the species list detected by eDNA analyses with environmental data of each survey site. Regarding aquatic insects, the species that inhabit each watershed tend to differ as the environment changes from upstream to downstream (Gallardo- Mayenco et a., 1998; Okamoto & Tojo, 2021; Okamoto et al., 2022). As a result of cluster analyses for the community structures of aquatic insects using eDNA data, we observed that the community structures of the aquatic insects differ greatly between the sites with upstream characteristics (St. 2, 3, 4, 6) and those with downstream characteristics (St. 1 and 5). In the upstream environments, St. 2 and 6 exhibited the most similar community structures. These sites had a higher altitude and riverbed slope degree than St. 3 and 4. In other words, we confirmed that the community structure is influenced by the river environment.

Since eDNA flows downstream, it may be expected that the species list detected from eDNA analyses at one site might be reflected in a community structure from further upstream. However, our results suggest that the community structure is more consistent with the environment of the site where the water was sampled. It appears that the range over which effluent eDNA is detectable may not be so extensive. According to a previous study on eDNA diffusion of fish, the eDNA only drifted 2 km downstream, at the farthest (Jo & Yamanaka, 2022). Therefore, the community structure detected by eDNA analyses reflects a relatively narrow range immediately above the sampling site. In the future, it is necessary to investigate the geographical scale and range of detectable eDNA in more detail.

## Conclusion

A previous study showed that it is difficult to design universal primers for detection in the mtDNA COI region, which is considered the general metabarcoding region of insects, because there is no continuous and stable region for comparing insect mitochondrial genomes registered in GenBank (Takenaka et al., 2023). In the present study, eDNA analyses were conducted based on the mtDNA 16S rRNA region, not the mtDNA COI region. Accurate eDNA analyses were demonstrated, while greatly expanding the database of aquatic insects in Kanagawa Prefecture, the target area. Notably, there is no such database for other regions of Japanese Islands, or elsewhere in the world. Furthermore, especially for insects with high genetic diversity, it is necessary to create a database specific to each region. In general, eDNA analyses use a 97% match for identification. Therefore, even if the same species is present in a different region, it may not be identified as a match. Although the mtDNA 16S rRNA region has fewer polymorphisms than the mtDNA COI region (Marquina et al., 2019), there are large genetic distances between each region, even within the same species, even in the mtDNA 16S rRNA region (Takenaka et al., 2019; Tojo et al., 2021). Nevertheless, eDNA analysis is an essential tool for performing environmental and biodiversity assessments. It is important to note that 8 species could not be distinguished by eDNA analysis, and thus they were treated as one taxonomic group in this study. Regarding the taxa that cannot be amplified, it is necessary to investigate whether they cannot be distinguished by the DNA barcode region when amplified using the MtInsects-16S primers set, or whether they cannot be distinguished by molecular markers.

As a major issue for the future, databases must be improved in each region. We showed that highly accurate results can be achieved when a regionally specific database is created. However, some important taxonomic groups remain unaccounted for, such as Diptera, because of the limited number of registered sequences in the Diptera database, and thus we could not conduct effective analyses. Eventually, this method will become a strong tool by expanding the databases for each region. Finally, we recommend the mtDNA 16S rRNA region as the standard DNA barcoding region for insects.

## Acknowledgments

We express our thanks to Dr. Suzuki T. (Kyoto University) for their valuable advice and encouragement. We are sincerely grateful to Bioengineering Lab Co., Ltd. and Plantbio Co., Ltd. for constructing the DNA database for insects. This study was supported by the River Fund (2021-5311- 005, 2022-5311-016 to MT) of the River Foundation, by a research grant from the Institute of Mountain Science, Shinshu University (MT).

