## Supplemental_file_MT for "Definitive environmental DNA research on aquatic insects: Analysis optimization using the recently developed MtInsects-16S primers set"

*, The contributions of the first and second authors are equal

**, Correspondence: Masaki Takenaka

Department of Biology, Faculty of Science, Shinshu University, Asahi 3-1-1, Matsumoto, Nagano 390-8621, Japan

**Supporting/Supplemental information**

Table S1. The count of reads within all reads detected by eDNA analysis in major animal taxa

| | | St. 1 | | | | | | St. 2 | | | | | |
|---|---|---|---|---|---|---|---|---|---|---|---|---|---|
| (sub)phylum | taxa | I | II | III | IV | V | VI | I | II | III | IV | V | VI |
| Arthropoda | Insecta | 107576 | 119252 | 138177 | 117889 | 136672 | 115425 | 206096 | 135770 | 187655 | 122966 | 143367 | 110868 |
| Arthropoda | Hexapoda | 286 | 1561 | 566 | 2206 | 652 | 1781 | 102 | 113 | 13 | 81 | 22 | 36 |
| Crustacea | Malacostraca | 116 | 85 | 141 | 102 | 114 | 98 | 51 | 53 | 60 | 52 | 31 | 35 |
| Crustacea | Oligostraca | 0 | 0 | 0 | 0 | 0 | 0 | 0 | 0 | 0 | 0 | 0 | 0 |
| Crustacea | Branchiopoda | 2280 | 4147 | 3392 | 5369 | 3272 | 4647 | 0 | 0 | 0 | 0 | 0 | 0 |
| Crustacea | Ostracoda | 0 | 14 | 0 | 0 | 0 | 29 | 12 | 0 | 0 | 0 | 0 | 0 |
| Myriapoda | Diplopoda | 0 | 0 | 0 | 0 | 0 | 0 | 0 | 16 | 0 | 0 | 0 | 0 |
| Myriapoda | Chilopoda | 42 | 17 | 24 | 42 | 24 | 72 | 0 | 0 | 0 | 41 | 0 | 34 |
| Chelicerata | Arachnida | 13 | 0 | 0 | 0 | 77 | 0 | 8 | 0 | 0 | 13 | 0 | 0 |
| Annelida | Oligochaeta | 9 | 78 | 0 | 0 | 32 | 120 | 0 | 0 | 0 | 36 | 0 | 20 |
| Mollusca | Gastropoda | 25 | 80 | 21 | 76 | 35 | 50 | 0 | 0 | 0 | 0 | 0 | 0 |
| Mollusca | Bivalvia | 0 | 11 | 9 | 42 | 17 | 0 | 0 | 0 | 0 | 0 | 0 | 0 |
| Cnidaria | Hydrozoa | 7180 | 30005 | 1598 | 7541 | 2486 | 6429 | 225 | 777 | 378 | 1011 | 254 | 886 |
| Platyhelminthes | Catenulida | 25 | 0 | 18 | 0 | 0 | 0 | 0 | 0 | 11 | 14 | 606 | 872 |
| Platyhelminthes | Rhabditophora | 0 | 0 | 0 | 0 | 0 | 0 | 0 | 0 | 0 | 0 | 0 | 0 |
| Nemertea | Enopla | 0 | 38 | 12 | 14 | 0 | 77 | 0 | 0 | 0 | 0 | 0 | 0 |
| Bryozoa | Gymnolaemata | 171 | 279 | 235 | 271 | 407 | 435 | 0 | 0 | 0 | 0 | 0 | 0 |
| Bryozoa | Phylactolaemata | 26 | 231 | 63 | 133 | 42 | 164 | 0 | 0 | 0 | 0 | 0 | 0 |

| | | | | | | | | | | | | | |
|---|---|---|---|---|---|---|---|---|---|---|---|---|---|
| Rotifera | Monogononta | 0 | 0 | 0 | 0 | 0 | 0 | 0 | 0 | 0 | 0 | 0 | 0 |
| Porifera | Demospongiae | 0 | 9 | 0 | 0 | 0 | 10 | 0 | 0 | 0 | 0 | 0 | 0 |
| Chordata | Mammalia | 86 | 578 | 118 | 1484 | 153 | 1022 | 37 | 36 | 37 | 34 | 0 | 0 |
| Chordata | Aves | 0 | 0 | 0 | 0 | 0 | 0 | 0 | 131 | 0 | 133 | 9 | 154 |
| Chordata | Amphibia | 0 | 36 | 0 | 51 | 0 | 29 | 0 | 105 | 20 | 126 | 0 | 126 |
| Chordata | Osteichthyes | 1719 | 6907 | 2265 | 8387 | 2855 | 9111 | 180 | 548 | 151 | 512 | 143 | 465 |
| Gastrotricha | Chaetonotida | 0 | 0 | 13 | 0 | 8 | 0 | 0 | 0 | 15 | 0 | 0 | 0 |
| Chlorophyta | Ulvophyceae | 0 | 0 | 0 | 0 | 9 | 0 | 0 | 0 | 0 | 0 | 0 | 0 |
| | Bacteria | 0 | 0 | 0 | 0 | 0 | 0 | 0 | 9 | 0 | 0 | 0 | 0 |
| | Viruses | 0 | 46 | | 17 | 0 | 40 | 0 | 0 | 0 | 0 | 0 | 0 |
| | other | 0 | 46 | 13 | 17 | 17 | 40 | 0 | 9 | 15 | 0 | 0 | 0 |
| | No Align | 12271 | 6314 | 13412 | 6273 | 16022 | 6544 | 3815 | 2666 | 6442 | 4066 | 2989 | 3416 |

Table S1 continued

| | | St. 3 | | | | | | St. 4 | | | | | |
|---|---|---|---|---|---|---|---|---|---|---|---|---|---|
| (sub)phylum | taxa | I | II | III | IV | V | VI | I | II | III | IV | V | VI |
| Arthropoda | Insecta | 198938 | 116379 | 180887 | 129624 | 158712 | 168699 | 90245 | 114762 | 192314 | 132942 | 172969 | 141040 |
| Arthropoda | Hexapoda | 210 | 452 | 196 | 287 | 136 | 591 | 0 | 0 | 10 | 0 | 0 | 19 |
| Crustacea | Malacostraca | 27 | 24 | 0 | 0 | 35 | 19 | 0 | 0 | 0 | 0 | 36 | 78 |
| Crustacea | Oligostraca | 0 | 0 | 0 | 0 | 0 | 0 | 0 | 0 | 0 | 0 | 0 | 0 |

| | | | | | | | | | | | | | |
|---|---|---|---|---|---|---|---|---|---|---|---|---|---|
| Crustacea | Branchiopoda | 0 | 0 | 0 | 0 | 0 | 0 | 0 | 0 | 0 | 0 | 0 | 0 |
| Crustacea | Ostracoda | 13 | 0 | 0 | 0 | 10 | 0 | 0 | 0 | 0 | 0 | 25 | 0 |
| Myriapoda | Diplopoda | 0 | 0 | 0 | 0 | 0 | 0 | 0 | 0 | 0 | 0 | 0 | 0 |
| Myriapoda | Chilopoda | 0 | 0 | 0 | 0 | 0 | 0 | 0 | 0 | 0 | 0 | 0 | 0 |
| Chelicerata | Arachnida | 0 | 0 | 0 | 0 | 0 | 0 | 0 | 0 | 0 | 0 | 0 | 0 |
| Annelida | Oligochaeta | 10 | 55 | 0 | 14 | 0 | 51 | 0 | 0 | 0 | 0 | 0 | 0 |
| Mollusca | Gastropoda | 8 | 25 | 19 | 15 | 10 | 10 | 0 | 0 | 0 | 0 | 0 | 0 |
| Mollusca | Bivalvia | 0 | 0 | 0 | 0 | 0 | 0 | 0 | 0 | 0 | 0 | 0 | 0 |
| Cnidaria | Hydrozoa | 203 | 643 | 231 | 534 | 158 | 757 | 0 | 26 | 0 | 23 | 0 | 45 |
| Platyhelminthes | Catenulida | 0 | 0 | 0 | 0 | 0 | 32 | 0 | 0 | 0 | 0 | 0 | 0 |
| Platyhelminthes | Rhabditophora | 0 | 0 | 0 | 0 | 0 | 0 | 0 | 0 | 0 | 0 | 0 | 0 |
| Nemertea | Enopla | 0 | 0 | 0 | 0 | 0 | 0 | 0 | 0 | 0 | 0 | 0 | 0 |
| Bryozoa | Gymnolaemata | 0 | 0 | 0 | 0 | 0 | 0 | 0 | 0 | 0 | 0 | 0 | 0 |
| Bryozoa | Phylactolaemata | 0 | 0 | 0 | 0 | 0 | 0 | 0 | 0 | 0 | 0 | 0 | 0 |
| Rotifera | Monogononta | 0 | 0 | 0 | 0 | 0 | 0 | 0 | 0 | 0 | 0 | 0 | 0 |
| Porifera | Demospongiae | 0 | 0 | 0 | 0 | 0 | 0 | 0 | 0 | 0 | 0 | 0 | 0 |
| Chordata | Mammalia | 0 | 34 | 0 | 0 | 0 | 24 | 10 | 0 | 13 | 28 | 0 | 37 |
| Chordata | Aves | 0 | 17 | 0 | 0 | 0 | 11 | 0 | 0 | 0 | 0 | 0 | 0 |
| Chordata | Amphibia | 8 | 0 | 0 | 10 | 0 | 12 | 0 | 0 | 0 | 27 | 13 | 54 |
| Chordata | Osteichthyes | 245 | 663 | 114 | 329 | 168 | 608 | 0 | 15 | 28 | 34 | 0 | 55 |
| Gastrotricha | Chaetonotida | 0 | 0 | 0 | 0 | 0 | 0 | 0 | 0 | 0 | 0 | 0 | 0 |
| Chlorophyta | Ulvophyceae | 0 | 0 | 0 | 0 | 0 | 36 | 0 | 0 | 0 | 0 | 0 | 0 |

| | | | | | | | | | | | | | |
|---|---|---|---|---|---|---|---|---|---|---|---|---|---|
| | Bacteria | 0 | 0 | 0 | 0 | 0 | 0 | 0 | 0 | 0 | 0 | 0 | 0 |
| | Viruses | 0 | 0 | 0 | 0 | 0 | 0 | 0 | 0 | 0 | 0 | 0 | 0 |
| | other | 0 | 0 | 0 | 0 | 0 | 11 | 0 | 0 | 0 | 0 | 0 | 36 |
| | No Align | 3735 | 921 | 2726 | 451 | 3155 | 1781 | 156 | 165 | 879 | 216 | 1264 | 2214 |

Table S1 continued

| | | St. 5 | | | | | | St. 6 | | | | | |
|---|---|---|---|---|---|---|---|---|---|---|---|---|---|
| (sub)phylum | taxa | I | II | III | IV | V | VI | I | II | III | IV | V | VI |
| Arthropoda | Insecta | 65946 | 93235 | 100521 | 76848 | 110694 | 74711 | 85207 | 129892 | 74056 | 139504 | 76291 | 89484 |
| Arthropoda | Hexapoda | 0 | 0 | 18 | 17 | 0 | 0 | 0 | 37 | 0 | 120 | 10 | 24 |
| Crustacea | Malacostraca | 0 | 0 | 0 | 0 | 0 | 0 | 0 | 0 | 0 | 0 | 0 | 0 |
| Crustacea | Oligostraca | 16 | 0 | 33 | 0 | 0 | 0 | 0 | 0 | 0 | 0 | 0 | 0 |
| Crustacea | Branchiopoda | 127 | 247 | 97 | 183 | 356 | 397 | 0 | 0 | 0 | 0 | 0 | 0 |
| Crustacea | Ostracoda | 0 | 0 | 0 | 0 | 0 | 0 | 55 | 0 | 0 | 0 | 0 | 0 |
| Myriapoda | Diplopoda | 0 | 0 | 0 | 0 | 0 | 0 | 0 | 0 | 0 | 0 | 0 | 0 |
| Myriapoda | Chilopoda | 62 | 70 | 0 | 8 | 8 | 36 | 13 | 0 | 0 | 225 | 0 | 27 |
| Chelicerata | Arachnida | 0 | 0 | 0 | 0 | 0 | 0 | 0 | 0 | 0 | 0 | 0 | 0 |
| Annelida | Oligochaeta | 0 | 0 | 0 | 0 | 0 | 0 | 0 | 21 | 0 | 0 | 0 | 0 |
| Mollusca | Gastropoda | 0 | 0 | 0 | 0 | 0 | 0 | 0 | 0 | 0 | 0 | 0 | 0 |

| | | | | | | | | | | | | | |
|---|---|---|---|---|---|---|---|---|---|---|---|---|---|
| Mollusca | Bivalvia | 0 | 0 | 0 | 0 | 0 | 0 | 0 | 0 | 0 | 0 | 0 | 0 |
| Cnidaria | Hydrozoa | 107 | 726 | 43 | 761 | 326 | 1219 | 25 | 135 | 94 | 47 | 47 | 53 |
| Platyhelminthes | Catenulida | 0 | 0 | 0 | 0 | 0 | 0 | 0 | 0 | 0 | 10 | 0 | 17 |
| Platyhelminthes | Rhabditophora | 0 | 0 | 0 | 0 | 0 | 0 | 0 | 0 | 0 | 0 | 0 | 0 |
| Nemertea | Enopla | 0 | 0 | 0 | 0 | 0 | 0 | 0 | 0 | 0 | 0 | 0 | 10 |
| Bryozoa | Gymnolaemata | 2477 | 3954 | 3233 | 2732 | 3288 | 2449 | 0 | 0 | 0 | 0 | 0 | 0 |
| Bryozoa | Phylactolaemata | 0 | 0 | 0 | 0 | 0 | 0 | 0 | 0 | 0 | 0 | 0 | 0 |
| Rotifera | Monogononta | 0 | 0 | 0 | 0 | 0 | 0 | 0 | 0 | 0 | 0 | 0 | 0 |
| Porifera | Demospongiae | 0 | 0 | 0 | 0 | 0 | 0 | 0 | 0 | 0 | 0 | 0 | 0 |
| Chordata | Mammalia | 0 | 0 | 19 | 30 | 0 | 0 | 0 | 33 | 172 | 4966 | 0 | 0 |
| Chordata | Aves | 0 | 0 | 0 | 0 | 0 | 0 | 0 | 11 | 0 | 0 | 0 | 0 |
| Chordata | Amphibia | 8 | 0 | 0 | 0 | 0 | 0 | 0 | 339 | 23 | 492 | 9 | 156 |
| Chordata | Osteichthyes | 9 | 307 | 45 | 486 | 138 | 655 | 0 | 44 | 0 | 106 | 21 | 33 |
| Gastrotricha | Chaetonotida | 0 | 0 | 0 | 0 | 0 | 0 | 0 | 0 | 0 | 0 | 0 | 0 |
| Chlorophyta | Ulvophyceae | 0 | 0 | 0 | 0 | 0 | 0 | 52 | 125 | 25 | 310 | 30 | 75 |
| | Bacteria | 0 | 0 | 0 | 0 | 0 | 11 | 0 | 0 | 0 | 46 | 0 | 0 |
| | Viruses | 0 | 0 | 0 | 0 | 0 | 0 | 0 | 0 | 0 | 0 | 0 | 0 |
| | other | 0 | 0 | 0 | 0 | 0 | 0 | 52 | 125 | 25 | 356 | 30 | 75 |
| | No Align | 89 | 23 | 215 | 27 | 384 | 88 | 901 | 3038 | 750 | 3209 | 862 | 2100 |

Table S2. The count of reads detected by eDNA analysis at order level within Insecta

| | St. 1 | | | | | | St. 2 | | | | | |
|---|---|---|---|---|---|---|---|---|---|---|---|---|
| | I | II | III | IV | V | VI | I | II | III | IV | V | VI |
| Ephemeroptera | 47745 | 51204 | 62663 | 44134 | 53561 | 36484 | 102629 | 46016 | 92910 | 52131 | 72909 | 45705 |
| Odonata | 316 | 457 | 550 | 363 | 454 | 413 | 382 | 216 | 251 | 239 | 180 | 246 |
| Plecoptera | 209 | 52 | 231 | 304 | 269 | 225 | 18356 | 7667 | 13981 | 8618 | 11684 | 7495 |
| Dermaptera | 0 | 0 | 33 | 34 | 37 | 65 | 0 | 95 | 63 | 128 | 25 | 0 |
| Orthoptera | 119 | 262 | 476 | 483 | 359 | 267 | 71 | 89 | 112 | 122 | 0 | 50 |
| Phasmatodea | 0 | 40 | 0 | 0 | 0 | 0 | 0 | 0 | 0 | 62 | 0 | 0 |
| Mantodea | 0 | 0 | 0 | 52 | 78 | 0 | 0 | 0 | 0 | 0 | 0 | 0 |
| Blattodea | 0 | 0 | 66 | 0 | 0 | 0 | 27563 | 10373 | 27334 | 11985 | 24303 | 12595 |
| Thysanoptera | 0 | 0 | 0 | 0 | 0 | 0 | 1088 | 19003 | 0 | 0 | 0 | 0 |
| Hemiptera | 1467 | 1005 | 1709 | 1307 | 2230 | 1509 | 1765 | 1297 | 1398 | 1238 | 1172 | 1323 |
| Psocodea | 67 | 183 | 128 | 193 | 195 | 171 | 5475 | 4309 | 5457 | 4048 | 3227 | 4407 |
| Hymenoptera | 2063 | 2254 | 1606 | 1487 | 1876 | 1508 | 2457 | 2096 | 2431 | 1639 | 1503 | 1855 |
| Raphidioptera | 0 | 0 | 0 | 0 | 0 | 0 | 0 | 0 | 0 | 0 | 0 | 0 |
| Megaloptera | 59 | 34 | 39 | 47 | 0 | 47 | 0 | 104 | 244 | 74 | 92 | 59 |
| Neuroptera | 0 | 0 | 65 | 27 | 0 | 0 | 233 | 345 | 813 | 700 | 567 | 574 |
| Coleoptera | 4244 | 4816 | 5287 | 5694 | 7195 | 5417 | 3149 | 2010 | 3723 | 2457 | 2058 | 2347 |
| Mecoptera | 0 | 0 | 0 | 0 | 0 | 0 | 0 | 0 | 0 | 0 | 0 | 0 |
| Diptera | 14534 | 16847 | 21731 | 18948 | 17688 | 16689 | 20675 | 15876 | 17696 | 14944 | 11506 | 12009 |
| Lepidoptera | 3890 | 4940 | 4607 | 4513 | 5291 | 4791 | 3463 | 2932 | 2683 | 2991 | 2140 | 2423 |

| | | | | | | | | | | | | |
|---|---|---|---|---|---|---|---|---|---|---|---|---|
| Trichoptera | 17087 | 18049 | 25211 | 25157 | 25132 | 20428 | 7532 | 7895 | 8372 | 9823 | 5376 | 9264 |
| All | 91800 | 100143 | 124402 | 102743 | 114365 | 88014 | 194838 | 120323 | 177468 | 111199 | 136742 | 100352 |

Table S2 continued

| | St. 3 | | | | | | St. 4 | | | | | |
|---|---|---|---|---|---|---|---|---|---|---|---|---|
| | I | II | III | IV | V | VI | I | II | III | IV | V | VI |
| Ephemeroptera | 57130 | 25658 | 46488 | 27496 | 54036 | 45011 | 77029 | 92669 | 69850 | 45401 | 98858 | 76977 |
| Odonata | 57 | 23 | 16 | 0 | 0 | 36 | 0 | 0 | 0 | 0 | 0 | 0 |
| Plecoptera | 1278 | 739 | 924 | 513 | 1342 | 1176 | 361 | 556 | 1877 | 1046 | 3798 | 1912 |
| Dermaptera | 0 | 0 | 0 | 0 | 0 | 0 | 0 | 0 | 0 | 0 | 0 | 0 |
| Orthoptera | 96 | 53 | 0 | 12 | 9 | 158 | 30 | 12 | 16 | 19 | 0 | 50 |
| Phasmatodea | 40 | 9 | 0 | 0 | 9 | 0 | 0 | 0 | 0 | 0 | 0 | 0 |
| Mantodea | 0 | 15 | 0 | 0 | 0 | 16 | 0 | 0 | 0 | 0 | 0 | 0 |
| Blattodea | 0 | 0 | 0 | 0 | 10 | 0 | 0 | 0 | 0 | 0 | 0 | 9 |
| Thysanoptera | 0 | 0 | 0 | 0 | 0 | 0 | 0 | 0 | 0 | 0 | 0 | 0 |
| Hemiptera | 2012 | 1154 | 473 | 218 | 1083 | 1509 | 0 | 52 | 166 | 156 | 153 | 363 |
| Psocodea | 492 | 476 | 201 | 299 | 423 | 666 | 13 | 66 | 129 | 129 | 153 | 151 |
| Hymenoptera | 1080 | 721 | 568 | 505 | 672 | 863 | 24 | 42 | 291 | 129 | 225 | 222 |
| Raphidioptera | 17 | 0 | 0 | 0 | 0 | 0 | 0 | 0 | 0 | 0 | 0 | 0 |

| | | | | | | | | | | | | |
|---|---|---|---|---|---|---|---|---|---|---|---|---|
| Megaloptera | 11 | 0 | 0 | 0 | 0 | 0 | 0 | 0 | 0 | 0 | 0 | 0 |
| Neuroptera | 112 | 30 | 31 | 25 | 0 | 117 | 0 | 0 | 0 | 0 | 0 | 0 |
| Coleoptera | 3085 | 1909 | 1289 | 1081 | 1898 | 2595 | 274 | 397 | 1461 | 1154 | 753 | 649 |
| Mecoptera | 0 | 0 | 0 | 0 | 0 | 0 | 0 | 0 | 0 | 0 | 0 | 0 |
| Diptera | 110858 | 63342 | 98835 | 69393 | 82145 | 85824 | 9346 | 14132 | 109738 | 76544 | 57387 | 44917 |
| Lepidoptera | 1150 | 900 | 559 | 575 | 1443 | 1467 | 9 | 13 | 114 | 34 | 225 | 198 |
| Trichoptera | 12180 | 11770 | 5683 | 6047 | 8646 | 15636 | 1278 | 2631 | 5950 | 5058 | 7016 | 9299 |
| All | 189598 | 106799 | 155067 | 106164 | 151716 | 155074 | 88364 | 110570 | 189592 | 129670 | 168568 | 134747 |

Table S2 continued

| | St. 5 | | | | | | St. 6 | | | | | |
|---|---|---|---|---|---|---|---|---|---|---|---|---|
| | I | II | III | IV | V | VI | I | II | III | IV | V | VI |
| Ephemeroptera | 46194 | 57531 | 81791 | 53004 | 60486 | 35754 | 49043 | 62805 | 43289 | 66328 | 43113 | 43875 |
| Odonata | 0 | 80 | 18 | 35 | 0 | 0 | 0 | 0 | 0 | 0 | 0 | 0 |
| Plecoptera | 401 | 472 | 328 | 187 | 806 | 267 | 2656 | 3261 | 1518 | 2370 | 2073 | 1746 |
| Dermaptera | 0 | 0 | 0 | 0 | 0 | 0 | 0 | 79 | 0 | 0 | 0 | 80 |
| Orthoptera | 0 | 0 | 13 | 0 | 16 | 42 | 0 | 143 | 0 | 0 | 0 | 55 |
| Phasmatodea | 0 | 0 | 0 | 0 | 0 | 0 | 34 | 77 | 264 | 209 | 103 | 51 |
| Mantodea | 0 | 0 | 0 | 0 | 0 | 0 | 0 | 0 | 0 | 0 | 0 | 0 |

| | | | | | | | | | | | | |
|---|---|---|---|---|---|---|---|---|---|---|---|---|
| Blattodea | 0 | 0 | 0 | 0 | 0 | 0 | 412 | 834 | 403 | 395 | 627 | 304 |
| Thysanoptera | 0 | 0 | 0 | 0 | 0 | 0 | 0 | 14 | 0 | 21 | 0 | 0 |
| Hemiptera | 10 | 124 | 0 | 109 | 222 | 109 | 1978 | 2582 | 2243 | 3748 | 3061 | 2650 |
| Psocodea | 23 | 0 | 13 | 0 | 48 | 12 | 2633 | 7653 | 2177 | 7154 | 2382 | 3720 |
| Hymenoptera | 90 | 112 | 150 | 127 | 312 | 186 | 757 | 739 | 364 | 861 | 534 | 584 |
| Raphidioptera | 0 | 0 | 0 | 0 | 0 | 0 | 0 | 0 | 0 | 0 | 0 | 0 |
| Megaloptera | 0 | 8 | 0 | 0 | 0 | 0 | 0 | 0 | 0 | 0 | 0 | 0 |
| Neuroptera | 0 | 0 | 0 | 9 | 0 | 0 | 55 | 0 | 122 | 0 | 37 | 29 |
| Coleoptera | 633 | 464 | 359 | 464 | 940 | 435 | 880 | 1377 | 595 | 2548 | 1125 | 1228 |
| Mecoptera | 0 | 0 | 0 | 0 | 0 | 0 | 0 | 0 | 0 | 116 | 0 | 0 |
| Diptera | 15601 | 27556 | 15197 | 18580 | 35650 | 27700 | 15069 | 25120 | 13342 | 26293 | 13651 | 19672 |
| Lepidoptera | 304 | 504 | 120 | 301 | 629 | 656 | 3664 | 5268 | 2874 | 5056 | 2796 | 2454 |
| Trichoptera | 2592 | 6249 | 2198 | 3723 | 8273 | 8412 | 4779 | 12828 | 4203 | 16325 | 3965 | 8308 |
| All | 65848 | 93100 | 100187 | 76539 | 107382 | 73573 | 81960 | 122780 | 71394 | 131424 | 73467 | 84756 |

Table S3 Comparison of the number of species detected by eDNA analyses and actual capture surveys.

| Order | Species | Database | St. 1 Yase-gawa river | | | | | | | | St. 2 Yataro-gawa river | | | | | | | | St. 3 Shijyuhasse-gawa river | | | | | | | |
|---|---|---|---|---|---|---|---|---|---|---|---|---|---|---|---|---|---|---|---|---|---|---|---|---|---|---|
| | | | Capture | | | | eDNA | | | | Capture | | | | eDNA | | | | Capture | | | | eDNA | | | |
| | | | qual | quan | | | 2-step PCR | | 3-step PCR | | qual | quan | | | 2-step PCR | | 3-step PCR | | qual | quan | | | 2-step PCR | | 3-step PCR | |
| | | | | 1 | 2 | 3 | reads | COM | reads | COM | | 1 | 2 | 3 | reads | COM | reads | COM | | 1 | 2 | 3 | reads | COM | reads | COM |
| Ephemeroptera | *Potamanthus formosus* | KNGDB | 93 | | | 1 | 2260 | Both | 908 | Both | 1 | | | | 0 | Capt | 0 | Capt | | | | | 0 | | 0 | |
| Ephemeroptera | *Acentrella gnom* | KNGDB | | | 1 | | 31 | Both | 58 | Both | | | | | 0 | | 0 | | | | | | 0 | | 16 | eDNA |
| Ephemeroptera | *Acentrella sibirica* | noDB | | 1 | | | 0 | noDB | 0 | noDB | | | | | 0 | | 0 | | 2 | 17 | | | 0 | noDB | 0 | noDB |
| Ephemeroptera | *Alainites yoshinensis* | KNGDB | | | | | 51 | eDNA | 91 | eDNA | 6 | 22 | 11 | 9 | 433 | Both | 879 | Both | 47 | 6 | 3 | 85 | 615 | Both | 1241 | Both |
| Ephemeroptera | *Baetiella japonica* | KNGDB | | | | | 660 | eDNA | 534 | eDNA | 2 | | | | 54 | Both | 169 | Both | 11 | 10 | 22 | 17 | 1566 | Both | 1776 | Both |
| Ephemeroptera | *Baetis sahoensis* | KNGDB | 8 | | | | 2004 | Both | 1755 | Both | | | | | 0 | | 19 | eDNA | 2 | | | | 2373 | Both | 2533 | Both |
| Ephemeroptera | *Baetis sp. F* | KNGDB | | | | | 0 | | 0 | | 1 | | | | 27 | Both | 37 | Both | | | | | 0 | | 0 | |
| Ephemeroptera | *Baetis sp. J* | KNGDB | | | | | 620 | eDNA | 389 | eDNA | | | | | 17 | eDNA | 0 | | 1 | 1 | 4 | 1 | 500 | Both | 395 | Both |
| Ephemeroptera | *Baetis taiwanensis* | KNGDB | 40 | | 19 | 37 | 45413 | Both | 36908 | Both | 1 | | | | 65 | Both | 0 | Capt | 14 | 4 | | | 10379 | Both | 9041 | Both |
| Ephemeroptera | *Baetis thermicus* | KNGDB | 15 | | | | 3236 | Both | 3495 | Both | 10 | 8 | 13 | 42 | 10379 | Both | 11928 | Both | 57 | 15 | 3 | 6 | 1916 | Both | 2837 | Both |
| Ephemeroptera | *Cloeon dipterum* | KNGDB | | | | | 24 | eDNA | 104 | eDNA | | | | | 0 | | 0 | | | | | | 0 | | 0 | |
| Ephemeroptera | *Labiobaetis atrebatinus orientalis* | KNGDB | 142 | | | | 0 | Capt | 0 | Capt | | | | | 0 | | 0 | | 8 | | | | 0 | Capt | 15 | Both |
| Ephemeroptera | *Nigrobaetis* | KNGDB | 2 | | | | 51 | Both | 122 | Both | | | | | 0 | | 0 | | | | | | 0 | | 0 | |

| | | | | | | | | | | | | | | | | | | | | | | | |
|---|---|---|---|---|---|---|---|---|---|---|---|---|---|---|---|---|---|---|---|---|---|---|---|
| | *acinaciger* | | | | | | | | | | | | | | | | | | | | | | |
| Ephemeroptera | *Nigrobaetis chocorata* | noDB | | | 2 | 4 | 0 | noDB | 0 | noDB | | | 0 | | 0 | | | | | 0 | | 0 | |
| Ephemeroptera | *Nigrobaetis latus* | KNGDB | | | | | 0 | | 0 | | 2 | | 422 | Both | 366 | Both | 3 | | | 0 | Capt | 15 | Both |
| Ephemeroptera | *Tenuibaetis flexifemora* | KNGDB | 15 | 34 | | 3 | 272 | Both | 827 | Both | | | 0 | | 0 | | | | | 0 | | 0 | |
| Ephemeroptera | *Tenuibaetis parvipterus* | ownDB | | | | | 10 | eDNA | 0 | | | | 31 | eDNA | 33 | eDNA | 3 | 1 | 1 | 299 | Both | 325 | Both |
| Ephemeroptera | *Ephoron shigae* | GenBank | 12 | 11 | 16 | 13 | 3024 | Both | 1631 | Both | | | 0 | | 0 | | | | | 0 | | 0 | |
| Ephemeroptera | *Isonychia valida* | KNGDB | | | | | 52 | eDNA | 8 | eDNA | | | 53 | eDNA | 0 | | | | | 0 | | 0 | |
| Ephemeroptera | *Choroterpes altioculus* | KNGDB | | | | | 0 | | 20 | eDNA | | | 0 | | 0 | | | | | 0 | | 0 | |
| Ephemeroptera | *Paraleptophlebia japonica* | GenBank | | | | | 0 | | 0 | | | | 2399 | eDNA | 2962 | eDNA | | | | 0 | | 0 | |
| Ephemeroptera | *Paraleptophlebia westoni* | KNGDB | | | | | 0 | | 0 | | | | 10 | eDNA | 0 | | | | | 0 | | 8 | eDNA |
| Ephemeroptera | *Bleptus fasciatus* | KNGDB | | | | | 0 | | 0 | | | | 109 | eDNA | 135 | eDNA | | | | 0 | | 0 | |
| Ephemeroptera | *Ecdyonurus kibunensis* | KNGDB | | | | | 0 | | 0 | | | 7 | 439 | Both | 234 | Both | | | | 0 | | 0 | |
| Ephemeroptera | *Ecdyonurus tobiironis* | KNGDB | | | | | 0 | | 0 | | | | 0 | | 0 | | | | | 0 | | 0 | |
| Ephemeroptera | *Ecdyonurus viridis* | KNGDB | | | | | 53 | eDNA | 0 | | 4 | | 16670 | Both | 10841 | Both | 5 | | | 2711 | Both | 1547 | Both |

| | | | | | | | | | | | | | | | | | | | | | | | | | | |
|---|---|---|---|---|---|---|---|---|---|---|---|---|---|---|---|---|---|---|---|---|---|---|---|---|---|---|
| Ephemeroptera | *Ecdyonurus yoshidae* | KNGDB | 1 | | | | 2387 | Both | 1270 | Both | | | | | 0 | | 0 | | | | | | 22 | eDNA | 28 | eDNA |
| Ephemeroptera | *Epeorus aesculus* | KNGDB | | | | | 0 | | 0 | | 6 | | 6 | 7 | 3401 | Both | 2261 | Both | | | | | 0 | | 0 | |
| Ephemeroptera | *Epeorus curvatulus* | KNGDB | | | | | 11 | eDNA | 0 | | 4 | 2 | | | 1543 | Both | 1781 | Both | 3 | | 1 | 19 | 431 | Both | 356 | Both |
| Ephemeroptera | *Epeorus latifolium / Epeorus l-nigrum / Epeorus napaeus* | KNGDB | 5 | 10 | 1 | 10 | 585 | Both | 536 | Both | 3 | 4 | 2 | 1 | 3205 | Both | 2777 | Both | 9 | 9 | | 1 | 144 | Both | 163 | Both |
| Ephemeroptera | *Epeorus nipponicus* | KNGDB | | | | | 0 | | 0 | | 14 | | | | 358 | Both | 402 | Both | | | | | 15 | eDNA | 18 | eDNA |
| Ephemeroptera | *Heptagenia kyotoensis* | KNGDB | | | | | 0 | | 0 | | | | | | 42 | eDNA | 17 | eDNA | | | | | 0 | | 0 | |
| Ephemeroptera | *Rhithrogena japonica* | KNGDB | | | | | 0 | | 0 | | | 6 | 19 | 37 | 13080 | Both | 8168 | Both | | | 7 | 1 | 5850 | Both | 3864 | Both |
| Ephemeroptera | *Rhithrogena tetrapunctigera* | KNGDB | | | | | 0 | | 0 | | | | 14 | 7 | 0 | Capt | 0 | Capt | | 1 | | | 0 | Capt | 0 | Capt |
| Ephemeroptera | *Cincticostella elongatula* | KNGDB | | | | | 0 | | 0 | | | | | | 202 | eDNA | 204 | eDNA | | | | | 117 | eDNA | 106 | eDNA |
| Ephemeroptera | *Cincticostella nigra* | KNGDB | | | | | 0 | | 0 | | 2 | | | 1 | 638 | Both | 491 | Both | 14 | | | | 524 | Both | 406 | Both |
| Ephemeroptera | *Drunella basalis* | KNGDB | | | | | 0 | | 0 | | | | | | 120 | eDNA | 13 | eDNA | | | | | 0 | | 0 | |
| Ephemeroptera | *Drunella* | noDB | | | | | 0 | | 0 | | | | | | 0 | | 0 | | | | | | 0 | | 0 | |

| | | | | | | | | | | | | | | | | | | | | | | | | | | |
|---|---|---|---|---|---|---|---|---|---|---|---|---|---|---|---|---|---|---|---|---|---|---|---|---|---|---|
| | *cryptomeria* | | | | | | | | | | | | | | | | | | | | | | | | | |
| Ephemeroptera | *Drunella ishiyamana* | KNGDB | | | | | 13 | eDNA | 0 | | 20 | 8 | 9 | 7 | 22 | Both | 20 | Both | 31 | 19 | 3 | 6 | 42 | Both | 37 | Both |
| Ephemeroptera | *Drunella sachalinensis / Drunella kohnoi* | KNGDB | | | | | 0 | | 0 | | 5 | | | 1 | 2274 | Both | 1578 | Both | 1 | | | | 572 | Both | 407 | Both |
| Ephemeroptera | *Drunella sp.1* | KNGDB | | | | | 0 | | 0 | | | | | | 219 | eDNA | 191 | eDNA | 16 | | | | 4474 | Both | 3352 | Both |
| Ephemeroptera | *Drunella trispina* | KNGDB | | | | | 0 | | 0 | | | | | 1 | 0 | Capt | 0 | Capt | | | | | 0 | | 0 | |
| Ephemeroptera | *Ephacerella longicaudata* | GenBank | | | | | 402 | eDNA | 399 | eDNA | | | | | 0 | | 0 | | | | | | 0 | | 0 | |
| Ephemeroptera | *Ephemerella aurivillii* | noDB | | | | | 0 | | 0 | | | | | | 0 | | 0 | | | 4 | | | 0 | noDB | 0 | noDB |
| Ephemeroptera | *Ephemerella ishiwatai* | noDB | | | | | 0 | | 0 | | | | | | 0 | | 0 | | | | | | 0 | | 0 | |
| Ephemeroptera | *Ephemerella occiprens* | KNGDB | 94 | | | | 1456 | Both | 1056 | Both | 1 | | | | 0 | Capt | 0 | Capt | 21 | | | | 120 | Both | 88 | Both |
| Ephemeroptera | *Ephemerella setigera* | KNGDB | | | | | 1379 | eDNA | 893 | eDNA | | | | | 138 | eDNA | 155 | eDNA | 7 | | | | 3354 | Both | 2519 | Both |
| Ephemeroptera | *Teleganopsis punctisetae* | KNGDB | 2 | 2 | 1 | 1 | 296 | Both | 76 | Both | | | | | 0 | | 0 | | 54 | | | 4 | 3360 | Both | 2281 | Both |
| Ephemeroptera | *Torleya japonica* | KNGDB | 6 | | 1 | 11 | 3861 | Both | 2817 | Both | | | | | 129 | eDNA | 118 | eDNA | | | | | 24 | eDNA | 0 | |
| Ephemeroptera | *Ephemera* | KNGDB | | | | | 166 | eDNA | 24 | eDNA | | | | | 0 | | 0 | | | | | | 0 | | 0 | |

| | | | | | | | | | | | | | | | | | | |
|---|---|---|---|---|---|---|---|---|---|---|---|---|---|---|---|---|---|---|
| | *orientalis* | | | | | | | | | | | | | | | | | |
| Ephemeroptera | *Ephemera strigata / Ephemera japonica* | KNGDB | 1058 | eDNA | 638 | eDNA | 45 | 1 | 2 | 1911 | Both | 1470 | Both | 5 | 216 | Both | 132 | Both |
| Plecoptera | *Kogotus sp.1* | KNGDB | 0 | | 0 | | | | | 0 | | 0 | | | 0 | | 10 | eDNA |
| Plecoptera | *Kogotus sp.2* | KNGDB | 0 | | 0 | | | | | 0 | | 0 | | | 0 | | 0 | |
| Plecoptera | *Pseudomegarcys japonica* | KNGDB | 0 | | 0 | | | | | 0 | | 17 | eDNA | | 0 | | 0 | |
| Plecoptera | *Amphinemura decemseta* | KNGDB | 55 | eDNA | 97 | eDNA | | | | 251 | eDNA | 154 | eDNA | | 19 | eDNA | 0 | |
| Plecoptera | *Amphinemura longispina* | KNGDB | 0 | | 0 | | | | | 310 | eDNA | 339 | eDNA | | 0 | | 0 | |
| Plecoptera | *Amphinemura zonata* | KNGDB | 0 | | 0 | | | | | 0 | | 0 | | | 0 | | 0 | |
| Plecoptera | *Indonemoura nohirae* | KNGDB | 0 | | 0 | | | | | 243 | eDNA | 121 | eDNA | | 0 | | 0 | |
| Plecoptera | *Nemoura chinonis* | KNGDB | 0 | | 0 | | | | | 26 | eDNA | 117 | eDNA | | 0 | | 0 | |
| Plecoptera | *Nemoura fulva / Nemoura redimiculum* | KNGDB | 0 | | 0 | | | | | 58 | eDNA | 56 | eDNA | | 0 | | 0 | |
| Plecoptera | *Nemoura* | KNGDB | 0 | | 0 | | | | | 0 | | 41 | eDNA | | 0 | | 0 | |

| | | | | | | | | | | | | | | | | | | |
|---|---|---|---|---|---|---|---|---|---|---|---|---|---|---|---|---|---|---|
| | *longicercia* | | | | | | | | | | | | | | | | | |
| Plecoptera | *Nemoura uenoi* | KNGDB | 0 | | 0 | | | | | | 120 | eDNA | 43 | eDNA | 8 | eDNA | 42 | eDNA |
| Plecoptera | *Protonemura orbiculata* | GenBank | 0 | | 0 | | | | | | 162 | eDNA | 28 | eDNA | 0 | | 0 | |
| Plecoptera | *Calineuria stigmatica* | KNGDB | 0 | | 0 | | | | | | 106 | eDNA | 143 | eDNA | 17 | eDNA | 0 | |
| Plecoptera | *Flavoperla hagiensis* | KNGDB | 0 | | 0 | | | | | | 757 | eDNA | 407 | eDNA | 0 | | 0 | |
| Plecoptera | *Flavoperla hatakeyamae* | KNGDB | 0 | | 0 | | | | | | 0 | | 0 | | 0 | | 0 | |
| Plecoptera | *Flavoperla thoracica* | KNGDB | 0 | | 0 | | | | | | 147 | eDNA | 333 | eDNA | 39 | eDNA | 49 | eDNA |
| Plecoptera | *Kamimuria quadrata* | KNGDB | 0 | | 0 | | 8 | 1 | 2 | 1 | 267 | Both | 180 | Both | 54 | eDNA | 11 | eDNA |
| Plecoptera | *Kamimuria tibialis* | KNGDB | 0 | | 0 | | | | | 1 | 0 | Capt | 0 | Capt | 0 | | 0 | |
| Plecoptera | *Kamimuria uenoi* | KNGDB | 0 | | 0 | | 7 | | 2 | 1 | 31 | Both | 65 | Both | 0 | | 0 | |
| Plecoptera | *Neoperla sp. 1 of Inada, 2011* | KNGDB | 8 | eDNA | 0 | | | | | | 0 | | 0 | | 0 | | 0 | |
| Plecoptera | *Neoperla sp. 3 of Inada, 2011* | KNGDB | 25 | eDNA | 83 | eDNA | | | | | 0 | | 0 | | 0 | | 0 | |
| Plecoptera | *Niponiella limbatella* | KNGDB | 0 | | 0 | | | | | | 0 | | 0 | | 0 | | 0 | |

| | | | | | | | | | | | | | | |
|---|---|---|---|---|---|---|---|---|---|---|---|---|---|---|
| Plecoptera | *Oyamia lugubris* | KNGDB | 0 | 0 | 1 | | 158 | Both | 77 | Both | 0 | | 0 | |
| Plecoptera | *Paragnetina suzukii* | KNGDB | 0 | 0 | | | 0 | | 0 | | 0 | | 0 | |
| Plecoptera | *Paragnetina tinctipennis* | KNGDB | 0 | 0 | 1 | | 88 | Both | 100 | Both | 0 | | 0 | |
| Plecoptera | *Cryptoperla japonica* | KNGDB | 0 | 0 | 8 | 1 | 1863 | Both | 1563 | Both | 0 | | 0 | |
| Plecoptera | *Paraleuctra okamotoa* | KNGDB | 0 | 0 | | | 418 | eDNA | 294 | eDNA | 0 | | 0 | |
| Plecoptera | *Perlomyia isobeae* | KNGDB | 0 | 0 | | | 28 | eDNA | 40 | eDNA | 0 | | 10 | eDNA |
| Plecoptera | *Rhopalopsole bulbifera* | GenBank | 0 | 0 | | | 317 | eDNA | 332 | eDNA | 0 | | 0 | |
| Plecoptera | *Rhopalopsole sinuacercia* | KNGDB | 0 | 0 | | | 52 | eDNA | 0 | | 0 | | 0 | |
| Plecoptera | *Alloperla nipponica* | KNGDB | 0 | 0 | | | 382 | eDNA | 153 | eDNA | 0 | | 0 | |
| Plecoptera | *Haploperla japonica* | KNGDB | 0 | 0 | | | 0 | | 0 | | 0 | | 0 | |
| Plecoptera | *Suwallia bimaculata* | GenBank | 0 | 0 | | | 2939 | eDNA | 2097 | eDNA | 963 | eDNA | 832 | eDNA |
| Plecoptera | *Sweltsa kibunensis* | KNGDB | 0 | 0 | | | 0 | | 38 | eDNA | 0 | | 0 | |
| Plecoptera | *Sweltsa nikkoensis* | KNGDB | 0 | 0 | | | 120 | eDNA | 82 | eDNA | 8 | eDNA | 0 | |

| | | | | | | | | | | | | | | | | |
|---|---|---|---|---|---|---|---|---|---|---|---|---|---|---|---|---|
| | */ Sweltsa kibunensis* | | | | | | | | | | | | | | | |
| Trichoptera | *Anisocentropus kawamurai* | KNGDB | 0 | | 25 | eDNA | 1 | 0 | Capt | 0 | Capt | | 0 | | 0 | |
| Trichoptera | *Nyctiophylax kisoensis* | KNGDB | 0 | | 0 | | | 52 | eDNA | 95 | eDNA | | 0 | | 0 | |
| Trichoptera | *Nothopsyche yamagataensis* | GenBank | 0 | | 0 | | 2 | 0 | Capt | 0 | Capt | | 0 | | 0 | |
| Trichoptera | *Eobrachycentrus vernalis* | KNGDB | 0 | | 0 | | | 0 | | 0 | | | 0 | | 0 | |
| Trichoptera | *Micrasema hanasense* | KNGDB | 0 | | 0 | | | 63 | eDNA | 101 | eDNA | 1 | 0 | Capt | 9 | Both |
| Trichoptera | *Micrasema uenoi* | KNGDB | 0 | | 0 | | 1 | 225 | Both | 481 | Both | | 0 | | 0 | |
| Trichoptera | *Lepidostoma japonicum* | KNGDB | 0 | | 52 | eDNA | | 267 | eDNA | 677 | eDNA | | 1934 | eDNA | 3302 | eDNA |
| Trichoptera | *Lepidostoma naraense* | KNGDB | 0 | | 0 | | | 0 | | 34 | eDNA | | 0 | | 0 | |
| Trichoptera | *Lepidostoma orientale* | KNGDB | 251 | eDNA | 279 | eDNA | | 0 | | 0 | | | 0 | | 0 | |
| Trichoptera | *Lepidostoma satoi* | KNGDB | 0 | | 0 | | | 0 | | 42 | eDNA | | 0 | | 0 | |
| Trichoptera | *Dolophilodes angustata* | KNGDB | 0 | | 0 | | | 12 | eDNA | 20 | eDNA | | 0 | | 0 | |

| | | | | | | | | | | | | | | | | | | | | | | | | | | | | | |
|---|---|---|---|---|---|---|---|---|---|---|---|---|---|---|---|---|---|---|---|---|---|---|---|---|---|---|---|---|---|
| Trichoptera | *Dolophilodes iroensis* | KNGDB | | | | | | 0 | | 0 | | | | | | | 17 | eDNA | 0 | | | | | | | 0 | | 0 | |
| Trichoptera | *Dolophilodes japonica* | KNGDB | | | | | | 0 | | 0 | | | 6 | | | | 738 | Both | 1558 | Both | | | | | | 0 | | 0 | |
| Trichoptera | *Kisaura minakawai* | KNGDB | | | | | | 0 | | 0 | | | | | | | 0 | | 0 | | | | | | | 25 | eDNA | 52 | eDNA |
| Trichoptera | *Kisaura tsudai* | KNGDB | | | | | | 0 | | 0 | | | | | | | 8 | eDNA | 39 | eDNA | | | | | | 0 | | 0 | |
| Trichoptera | *Apsilochorema sutshanum* | KNGDB | | | | | | 0 | | 0 | | | | | | 1 | 213 | Both | 288 | Both | | | | | | 0 | | 57 | eDNA |
| Trichoptera | *Psychomyia acutipennis* | KNGDB | | | | | | 24 | eDNA | 362 | eDNA | | | | | | 0 | | 0 | | | | | | | 0 | | 0 | |
| Trichoptera | *Neophylax japonicus* | KNGDB | | | | | | 0 | | 0 | | | 3 | 1 | 2 | 1 | 580 | Both | 952 | Both | | | | | | 0 | | 0 | |
| Trichoptera | *Uenoa tokunagai* | KNGDB | | | | | | 0 | | 0 | | | | 5 | 3 | 7 | 0 | Capt | 0 | Capt | | | | | | 0 | | 0 | |
| Trichoptera | *Gumaga orientalis* | KNGDB | | | | 30 | 9 | 0 | Capt | 0 | Capt | | | | | | 0 | | 0 | | | | | | | 0 | | 0 | |
| Trichoptera | *Apatania aberrans* | KNGDB | | | | | | 23 | eDNA | 22 | eDNA | | | | | | 24 | eDNA | 68 | eDNA | | | | | | 45 | eDNA | 106 | eDNA |
| Trichoptera | *Cheumatopsyche brevilineata* | KNGDB | | | | | | 2205 | eDNA | 1792 | eDNA | | | | | | 0 | | 0 | | | | | | | 0 | | 0 | |
| Trichoptera | *Cheumatopsyche infascia* | KNGDB | 31 | | 3 | 5 | 9 | 6218 | Both | 5521 | Both | | | | | | 0 | | 0 | | | | | | | 117 | eDNA | 126 | eDNA |
| Trichoptera | *Diplectrona kibuneana* | KNGDB | | | | | | 33 | eDNA | 77 | eDNA | | | | | | 63 | eDNA | 83 | eDNA | | | | | | 0 | | 0 | |

| | | | | | | | | | | | | | | | | | | | | | | | | | | |
|---|---|---|---|---|---|---|---|---|---|---|---|---|---|---|---|---|---|---|---|---|---|---|---|---|---|---|
| Trichoptera | *Hydropsyche albicephala* | KNGDB | | | | | 0 | | 0 | | 3 | 1 | 6 | 3 | 1449 | Both | 1808 | Both | | | | | 0 | | 0 | |
| Trichoptera | *Hydropsyche selysi / Hydropsyche gifuana* | KNGDB | | | | | 118 | eDNA | 101 | eDNA | | | | | 12 | eDNA | 0 | | | | | | 0 | | 0 | |
| Trichoptera | *Hydropsyche orientalis* | KNGDB | 41 | 176 | 8 | 90 | 3283 | Both | 3119 | Both | 5 | 2 | | 1 | 279 | Both | 491 | Both | 12 | 8 | 2 | 7 | 970 | Both | 1176 | Both |
| Trichoptera | *Hydropsyche setensis* | KNGDB | 1 | 25 | | 4 | 8863 | Both | 7491 | Both | | | | | 0 | | 0 | | | | | | 9 | eDNA | 0 | |
| Trichoptera | *Eubasilissa regina* | KNGDB | | | | | 0 | | 0 | | | | | | 12 | eDNA | 0 | | | | | | 0 | | 0 | |
| Trichoptera | *Himalopsyche japonica* | KNGDB | | | | | 0 | | 0 | | | | | | 0 | | 0 | | | | | | 0 | | 0 | |
| Trichoptera | *Rhyacophila brevicephala* | KNGDB | 1 | 4 | | | 22 | Both | 60 | Both | | | | 1 | 27 | Both | 14 | Both | 2 | | | 4 | 11 | Both | 0 | Capt |
| Trichoptera | *Rhyacophila clemens* | KNGDB | | | | | 0 | | 0 | | | | 2 | | 64 | Both | 114 | Both | | | | | 0 | | 0 | |
| Trichoptera | *Rhyacophila diffidens* | KNGDB | | | | | 0 | | 0 | | | | | | 19 | eDNA | 22 | eDNA | | | | | 0 | | 0 | |
| Trichoptera | *Rhyacophila kawamurae* | KNGDB | | | | | 17 | eDNA | 13 | eDNA | | | | 1 | 244 | Both | 358 | Both | | | | | 0 | | 8 | eDNA |
| Trichoptera | *Rhyacophila kisoensis* | KNGDB | | | | | 0 | | 0 | | | | | | 20 | eDNA | 0 | | | | | | 0 | | 0 | |

| | | | | | | | | | | | | | | | | | | | | | | | | | | |
|---|---|---|---|---|---|---|---|---|---|---|---|---|---|---|---|---|---|---|---|---|---|---|---|---|---|---|
| Trichoptera | *Rhyacophila towadensis / Rhyacophila lezeyi* | KNGDB | | | | | 0 | | 0 | | | | | | 24 | eDNA | 8 | eDNA | | | | | 0 | | 0 | |
| Trichoptera | *Rhyacophila nakagawai* | KNGDB | | | | | 0 | | 0 | | | | | | 353 | eDNA | 288 | eDNA | | | | | 12 | eDNA | 18 | eDNA |
| Trichoptera | *Rhyacophila nipponica / Rhyacophila nigrocephala* | KNGDB | | | | | 51 | eDNA | 16 | eDNA | | | 1 | | 67 | Both | 53 | Both | 8 | 1 | 1 | 4 | 171 | Both | 164 | Both |
| Trichoptera | *Rhyacophila shikotsuensis* | KNGDB | | | | | 0 | | 0 | | 7 | 11 | 4 | 4 | 11 | Both | 37 | Both | | | | | 0 | | 0 | |
| Trichoptera | *Rhyacophila yamanakensis* | KNGDB | | | | | 100 | eDNA | 87 | eDNA | | | | | 0 | | 0 | | | | | | 20 | eDNA | 0 | |
| Trichoptera | *Goera japonica* | KNGDB | 103 | | 10 | | 80 | Both | 96 | Both | | | | | 22 | eDNA | 66 | eDNA | 3 | 3 | | | 628 | Both | 1029 | Both |
| Trichoptera | *Stenopsyche marmorata* | KNGDB | 67 | 57 | 43 | 26 | 1996 | Both | 3291 | Both | | 1 | | | 92 | Both | 280 | Both | | | | | 25 | eDNA | 80 | eDNA |
| Trichoptera | *Mystacides azurea* | KNGDB | | | | | 0 | | 45 | eDNA | | | | | 0 | | 0 | | | | | | 0 | | 0 | |
| Trichoptera | *Hydroptila phenianica* | KNGDB | | | | | 10 | eDNA | 0 | | | | | | 0 | | 0 | | | | | | 0 | | 0 | |
| Trichoptera | *Oxyethira ecornuta* | GenBank | | | | | 0 | | 0 | | | | | | 0 | | 0 | | | | | | 0 | | 0 | |
| Trichoptera | *Phryganopsyche* | GenBank | | | | | 0 | | 0 | | | | | | 0 | | 0 | | | | | | 0 | | 0 | |

| | | | | | | | | | | | | | | | | | | | | | | | | | | |
|---|---|---|---|---|---|---|---|---|---|---|---|---|---|---|---|---|---|---|---|---|---|---|---|---|---|---|
| | *latipennis* | | | | | | | | | | | | | | | | | | | | | | | | | |
| Trichoptera | *Glossosoma ussuricum* | KNGDB | | | | | 98 | eDNA | 99 | eDNA | | | | | 1936 | eDNA | 4366 | eDNA | | | | | 2398 | eDNA | 5112 | eDNA |
| Trichoptera | *Glossosoma altaicum* | KNGDB | | | | | 0 | | 16 | eDNA | | | | | 0 | | 0 | | | | | | 411 | eDNA | 728 | eDNA |

Database, referenced database used for final species identification; ownDB, the DNA database for insects created by Kanagawa Prefecture. Capture, the number of insects collected by capture surveyt; qual, qualitative sampling; quan, quantitative sampling; eDNA, the number of DNA reads detected by eDNA analysis.

### Table S3 continued

| Order | Species | Database | St. 4 Nakatsu-gawa river | | | | | | | | St. 5 Sakawa-gawa river | | | | | | | | St. 6 Yoduku-gawa river | | | | | | | |
|---|---|---|---|---|---|---|---|---|---|---|---|---|---|---|---|---|---|---|---|---|---|---|---|---|---|---|
| | | | Capture | | | | eDNA | | | | Capture | | | | eDNA | | | | Capture | | | | eDNA | | | |
| | | | qual | quan | | | 2-step PCR | | 3-step PCR | | qual | quan | | | 2-step PCR | | 3-step PCR | | qual | quan | | | 2-step PCR | | 3-step PCR | |
| | | | | 1 | 2 | 3 | reads | COM | reads | COM | | 1 | 2 | 3 | reads | COM | reads | COM | | 1 | 2 | 3 | reads | COM | reads | COM |
| Ephemeroptera | Potamanthus formosus | KNGDB | | | | | 0 | | 0 | | | | | | 0 | | 0 | | 2 | | | | 0 | Capt | 0 | Capt |
| Ephemeroptera | Acentrella gnom | KNGDB | | | | | 0 | | 0 | | | | | | 0 | | 0 | | | | | | 0 | | 0 | |
| Ephemeroptera | Acentrella sibirica | noDB | | | | | 0 | | 0 | | | | | | 0 | | 0 | | | | | | 0 | | 0 | |
| Ephemeroptera | Alainites | KNGDB | 26 | 6 | 5 | | 1472 | Both | 3164 | Both | 23 | 8 | 6 | 13 | 45 | Both | 10 | Both | 12 | 2 | 2 | | 91 | Both | 89 | Both |

| | | | | | | | | | | | | | | | | | | | | | | | | | | |
|---|---|---|---|---|---|---|---|---|---|---|---|---|---|---|---|---|---|---|---|---|---|---|---|---|---|---|
| | yoshinensis | | | | | | | | | | | | | | | | | | | | | | | | | |
| Ephemeroptera | Baetiella japonica | KNGDB | 21 | 41 | 58 | 22 | 4291 | Both | 4048 | Both | 24 | 41 | 16 | 9 | 7032 | Both | 5822 | Both | 10 | 11 | 10 | 11 | 1604 | Both | 2041 | Both |
| Ephemeroptera | Baetis sahoensis | KNGDB | | | | | 15771 | eDNA | 15964 | eDNA | 34 | | | | 20413 | Both | 20295 | Both | | | | | 9332 | eDNA | 8039 | eDNA |
| Ephemeroptera | Baetis sp. F | KNGDB | 5 | | | | 0 | Capt | 0 | Capt | | | | | 0 | | 19 | eDNA | | | | | 0 | | 11 | eDNA |
| Ephemeroptera | Baetis sp. J | KNGDB | 3 | | | | 214 | Both | 287 | Both | 57 | 5 | 7 | 9 | 41006 | Both | 30733 | Both | 10 | | | | 0 | Capt | 0 | Capt |
| Ephemeroptera | Baetis taiwanensis | KNGDB | | 1 | | 1 | 51245 | Both | 47972 | Both | 18 | 1 | 6 | | 21597 | Both | 19951 | Both | 6 | | 4 | 3 | 0 | Capt | 161 | Both |
| Ephemeroptera | Baetis thermicus | KNGDB | 21 | 49 | 82 | 37 | 10798 | Both | 15847 | Both | 47 | | | 1 | 1403 | Both | 1487 | Both | 8 | 14 | 21 | 8 | 7735 | Both | 8563 | Both |
| Ephemeroptera | Cloeon dipterum | KNGDB | | | | | 0 | | 0 | | | | | | 0 | | 0 | | | | | | 0 | | 0 | |
| Ephemeroptera | Labiobaetis atrebatinus orientalis | KNGDB | | | | | 0 | | 0 | | 38 | | | | 0 | Capt | 0 | Capt | | | | | 0 | | 0 | |
| Ephemeroptera | Nigrobaetis acinaciger | KNGDB | | | | | 0 | | 0 | | 1 | | | | 49 | Both | 0 | Capt | | | | | 0 | | 0 | |
| Ephemeroptera | Nigrobaetis chocorata | noDB | | | | | 0 | | 0 | | | 1 | 2 | 3 | 0 | noDB | 0 | noDB | | | | | 0 | | 0 | |
| Ephemeroptera | Nigrobaetis latus | KNGDB | | | | | 17 | eDNA | 8 | eDNA | | | | | 0 | | 0 | | | | | | 0 | | 0 | |
| Ephemeroptera | Tenuibaetis flexifemora | KNGDB | | | | | 0 | | 0 | | 35 | | | | 2727 | Both | 3218 | Both | | | | | 0 | | 0 | |
| Ephemeroptera | Tenuibaetis parvipterus | ownDB | 1 | | | | 556 | Both | 409 | Both | 6 | 5 | | | 181 | Both | 148 | Both | | | | | 56 | eDNA | 14 | eDNA |
| Ephemeroptera | Ephoron shigae | GenBank | | | | | 0 | | 0 | | | | | | 0 | | 0 | | | | | | 0 | | 0 | |
| Ephemeroptera | Isonychia valida | KNGDB | | | | | 0 | | 0 | | 3 | | | | 277 | Both | 458 | Both | | | | | 0 | | 0 | |

| | | | | | | | | | | | | | | | | | | | | | | | | | | |
|---|---|---|---|---|---|---|---|---|---|---|---|---|---|---|---|---|---|---|---|---|---|---|---|---|---|---|
| Ephemeroptera | Choroterpes altioculus | KNGDB | | | | | 0 | | 0 | | | 1 | | | 0 | Capt | 0 | Capt | | | | | 0 | | 0 | |
| Ephemeroptera | Paraleptophlebia japonica | GenBank | | | | | 51 | eDNA | 13 | eDNA | | | | | 0 | | 0 | | | | | | 0 | | 0 | |
| Ephemeroptera | Paraleptophlebia westoni | KNGDB | | | | | 0 | | 0 | | | | | | 0 | | 0 | | | | | | 0 | | 0 | |
| Ephemeroptera | Bleptus fasciatus | KNGDB | | | | | 0 | | 0 | | | | | | 0 | | 0 | | | | | | 0 | | 0 | |
| Ephemeroptera | Ecdyonurus kibunensis | KNGDB | 40 | | | | 0 | Capt | 0 | Capt | | | | | 0 | | 0 | | 36 | | | | 205 | Both | 0 | Capt |
| Ephemeroptera | Ecdyonurus tobiironis | KNGDB | 3 | | | | 0 | Capt | 0 | Capt | | | | | 0 | | 0 | | 1 | | | | 0 | Capt | 0 | Capt |
| Ephemeroptera | Ecdyonurus viridis | KNGDB | 29 | | | | 1797 | Both | 1191 | Both | 31 | | | | 2112 | Both | 1776 | Both | 44 | | | | 3269 | Both | 1899 | Both |
| Ephemeroptera | Ecdyonurus yoshidae | KNGDB | 3 | | | | 0 | Capt | 0 | Capt | 27 | | | | 1579 | Both | 879 | Both | | | | | 0 | | 0 | |
| Ephemeroptera | Epeorus aesculus | KNGDB | | | | 2 | 75 | Both | 110 | Both | | | | | 0 | | 24 | eDNA | | | | | 384 | eDNA | 717 | eDNA |
| Ephemeroptera | Epeorus curvatulus | KNGDB | 12 | 1 | 3 | 1 | 616 | Both | 654 | Both | 8 | 3 | | 1 | 292 | Both | 285 | Both | 10 | 1 | | 6 | 3193 | Both | 2837 | Both |
| Ephemeroptera | Epeorus latifolium / Epeorus l-nigrum / Epeorus napaeus | KNGDB | 3 | | | 1 | 1069 | Both | 686 | Both | 11 | | 1 | 1 | 1603 | Both | 1671 | Both | 8 | | 3 | | 18976 | Both | 15842 | Both |
| Ephemeroptera | Epeorus nipponicus | KNGDB | | | | | 0 | | 26 | eDNA | 3 | | | | 61 | Both | 71 | Both | | | | | 0 | | 0 | |
| Ephemeroptera | Heptagenia | KNGDB | | | | | 0 | | 0 | | | | | | 0 | | 0 | | | | | | 0 | | 0 | |

| | | | | | | | | | | | | | | | | | | | | | | | | | | |
|---|---|---|---|---|---|---|---|---|---|---|---|---|---|---|---|---|---|---|---|---|---|---|---|---|---|---|
| | kyotoensis | | | | | | | | | | | | | | | | | | | | | | | | | |
| Ephemeroptera | Rhithrogena japonica | KNGDB | 12 | 41 | 46 | 32 | 17913 | Both | 11860 | Both | | | 1 | 1 | 828 | Both | 816 | Both | 3 | 3 | 6 | | 29933 | Both | 20831 | Both |
| Ephemeroptera | Rhithrogena tetrapunctigera | KNGDB | 4 | 10 | 26 | 2 | 0 | Capt | 0 | Capt | 2 | 3 | 8 | 2 | 312 | Both | 62 | Both | 2 | 2 | 3 | 2 | 0 | Capt | 0 | Capt |
| Ephemeroptera | Cincticostella elongatula | KNGDB | | | | | 48 | eDNA | 49 | eDNA | | | | | 0 | | 0 | | | | | | 0 | | 0 | |
| Ephemeroptera | Cincticostella nigra | KNGDB | 16 | 1 | | | 208 | Both | 173 | Both | | | | | 0 | | 0 | | | 2 | | | 84 | Both | 35 | Both |
| Ephemeroptera | Drunella basalis | KNGDB | | | | | 0 | | 0 | | | | | | 0 | | 0 | | | | | | 0 | | 0 | |
| Ephemeroptera | Drunella cryptomeria | noDB | | | 1 | | 0 | noDB | 0 | noDB | | | | | 0 | | 0 | | | | | | 0 | | 0 | |
| Ephemeroptera | Drunella ishiyamana | KNGDB | 58 | 21 | 44 | 19 | 0 | Capt | 0 | Capt | | | | | 437 | eDNA | 198 | eDNA | 4 | 3 | 6 | 4 | 46 | Both | 39 | Both |
| Ephemeroptera | Drunella sachalinensis / Drunella kohnoi | KNGDB | 30 | 4 | 2 | | 1174 | Both | 1075 | Both | | | | | 11 | eDNA | 0 | | 13 | 1 | | 2 | 2132 | Both | 1426 | Both |
| Ephemeroptera | Drunella sp.1 | KNGDB | | | | | 1052 | eDNA | 711 | eDNA | | | | | 16 | eDNA | 17 | eDNA | | | | | 191 | eDNA | 393 | eDNA |
| Ephemeroptera | Drunella trispina | KNGDB | | | | | 0 | | 0 | | | | | | 0 | | 0 | | 1 | | | | 0 | Capt | 0 | Capt |
| Ephemeroptera | Ephacerella longicaudata | GenBank | | | | | 0 | | 0 | | | | | | 0 | | 0 | | | | | | 0 | | 0 | |
| Ephemeroptera | Ephemerella | noDB | | | | | 0 | | 0 | | | | | | 0 | | 0 | | | | | | 0 | | 0 | |

| | | | | | | | | | | | | | | | | | | | | | | |
|---|---|---|---|---|---|---|---|---|---|---|---|---|---|---|---|---|---|---|---|---|---|---|
| | aurivillii | | | | | | | | | | | | | | | | | | | | | |
| Ephemeroptera | Ephemerella ishiwatai | noDB | 1 | | 0 | noDB | 0 | noDB | | | | | 0 | | 0 | | 2 | | 0 | noDB | 0 | noDB |
| Ephemeroptera | Ephemerella occiprens | KNGDB | | | 0 | | 0 | | 34 | | | | 935 | Both | 378 | Both | | | 0 | | 0 | |
| Ephemeroptera | Ephemerella setigera | KNGDB | | | 136 | eDNA | 155 | eDNA | 3 | | 1 | 1 | 3089 | Both | 2535 | Both | | | 922 | eDNA | 812 | eDNA |
| Ephemeroptera | Teleganopsis punctisetae | KNGDB | 8 | | 79 | Both | 56 | Both | 33 | 9 | 6 | 5 | 1030 | Both | 1171 | Both | | 1 | 0 | Capt | 12 | Both |
| Ephemeroptera | Torleya japonica | KNGDB | | 1 | 0 | Capt | 8 | Both | 6 | | | | 2325 | Both | 1738 | Both | | | 453 | eDNA | 215 | eDNA |
| Ephemeroptera | Ephemera orientalis | KNGDB | | | 0 | | 0 | | | | | | 0 | | 0 | | | | 0 | | 0 | |
| Ephemeroptera | Ephemera strigata / Ephemera japonica | KNGDB | 12 | | 304 | Both | 123 | Both | 3 | | | | 551 | Both | 231 | Both | 3 | | 1590 | Both | 1712 | Both |
| Plecoptera | Kogotus sp.1 | KNGDB | | | 0 | | 0 | | | | | | 0 | | 0 | | | | 0 | | 0 | |
| Plecoptera | Kogotus sp.2 | KNGDB | | | 0 | | 0 | | | | | | 0 | | 0 | | | | 343 | eDNA | 367 | eDNA |
| Plecoptera | Pseudomegarcys japonica | KNGDB | | | 0 | | 0 | | | | | | 0 | | 0 | | | | 43 | eDNA | 0 | |
| Plecoptera | Amphinemura decemseta | KNGDB | | | 0 | | 0 | | | | | | 0 | | 0 | | | | 0 | | 0 | |
| Plecoptera | Amphinemura longispina | KNGDB | | | 0 | | 0 | | | | | | 0 | | 0 | | | | 0 | | 0 | |

| | | | | | | | | | | | | | | | | |
|---|---|---|---|---|---|---|---|---|---|---|---|---|---|---|---|---|
| Plecoptera | Amphinemura zonata | KNGDB | | 0 | | 0 | | | 0 | | 0 | | 0 | | 0 | |
| Plecoptera | Indonemoura nohirae | KNGDB | | 17 | eDNA | 0 | | | 0 | | 0 | | 307 | eDNA | 291 | eDNA |
| Plecoptera | Nemoura chinonis | KNGDB | | 0 | | 0 | | | 0 | | 0 | | 0 | | 0 | |
| Plecoptera | Nemoura fulva / Nemoura redimiculum | KNGDB | | 8 | eDNA | 0 | | | 0 | | 0 | | 168 | eDNA | 107 | eDNA |
| Plecoptera | Nemoura longicercia | KNGDB | | 0 | | 0 | | | 0 | | 0 | | 40 | eDNA | 0 | |
| Plecoptera | Nemoura uenoi | KNGDB | | 0 | | 0 | | | 0 | | 0 | | 139 | eDNA | 113 | eDNA |
| Plecoptera | Protonemura orbiculata | GenBank | | 0 | | 0 | | | 0 | | 0 | | 0 | | 0 | |
| Plecoptera | Calineuria stigmatica | KNGDB | | 41 | eDNA | 50 | eDNA | | 0 | | 0 | | 479 | eDNA | 208 | eDNA |
| Plecoptera | Flavoperla hagiensis | KNGDB | | 11 | eDNA | 0 | | | 0 | | 0 | | 274 | eDNA | 139 | eDNA |
| Plecoptera | Flavoperla hatakeyamae | KNGDB | | 0 | | 0 | | | 48 | eDNA | 36 | eDNA | 0 | | 0 | |
| Plecoptera | Flavoperla thoracica | KNGDB | | 0 | | 0 | | | 0 | | 0 | | 149 | eDNA | 0 | |
| Plecoptera | Kamimuria | KNGDB | 1 | 0 | Capt | 0 | Capt | 1 | 8 | Both | 18 | Both | 0 | | 58 | eDNA |

| | | | | | | | | | | | | | | | | | |
|---|---|---|---|---|---|---|---|---|---|---|---|---|---|---|---|---|---|
| | quadrata | | | | | | | | | | | | | | | | |
| Plecoptera | Kamimuria tibialis | KNGDB | | 0 | | 0 | | 0 | | 0 | | | | 0 | | 0 | |
| Plecoptera | Kamimuria uenoi | KNGDB | | 0 | | 0 | | 0 | | 0 | | 4 | 1 | 0 | Capt | 0 | Capt |
| Plecoptera | Neoperla sp. 1 of Inada, 2011 | KNGDB | | 0 | | 0 | | 0 | | 0 | | | | 0 | | 0 | |
| Plecoptera | Neoperla sp. 3 of Inada, 2011 | KNGDB | | 0 | | 0 | | 192 | eDNA | 85 | eDNA | | | 0 | | 0 | |
| Plecoptera | Niponiella limbatella | KNGDB | | 0 | | 0 | | 0 | | 0 | | | | 0 | | 0 | |
| Plecoptera | Oyamia lugubris | KNGDB | | 0 | | 0 | | 0 | | 0 | | | | 0 | | 0 | |
| Plecoptera | Paragnetina suzukii | KNGDB | | 12 | eDNA | 0 | | 0 | | 0 | | | | 0 | | 0 | |
| Plecoptera | Paragnetina tinctipennis | KNGDB | | 0 | | 0 | | 0 | | 0 | | | | 0 | | 110 | eDNA |
| Plecoptera | Cryptoperla japonica | KNGDB | 1 | 0 | Capt | 10 | Both | 0 | | 0 | | | | 213 | eDNA | 108 | eDNA |
| Plecoptera | Paraleuctra okamotoa | KNGDB | | 0 | | 0 | | 0 | | 0 | | | | 0 | | 27 | eDNA |
| Plecoptera | Perlomyia isobeae | KNGDB | | 22 | eDNA | 17 | eDNA | 0 | | 0 | | | | 71 | eDNA | 0 | |
| Plecoptera | Rhopalopsole bulbifera | GenBank | | 0 | | 0 | | 0 | | 0 | | | | 54 | eDNA | 53 | eDNA |
| Plecoptera | Rhopalopsole | KNGDB | | 0 | | 0 | | 0 | | 0 | | | | 0 | | 0 | |

| | | | | | | | | | | | | | | | | | |
|---|---|---|---|---|---|---|---|---|---|---|---|---|---|---|---|---|---|
| | sinuacercia | | | | | | | | | | | | | | | | |
| Plecoptera | Alloperla nipponica | KNGDB | | 0 | | 0 | | | 0 | | 0 | | | 0 | | 0 | |
| Plecoptera | Haploperla japonica | KNGDB | | 0 | | 0 | | | 0 | | 0 | | | 0 | | 75 | eDNA |
| Plecoptera | Suwallia bimaculata | GenBank | | 1924 | eDNA | 1333 | eDNA | | 72 | eDNA | 98 | eDNA | | 1786 | eDNA | 1367 | eDNA |
| Plecoptera | Sweltsa kibunensis | KNGDB | | 0 | | 0 | | | 0 | | 0 | | | 0 | | 0 | |
| Plecoptera | Sweltsa nikkoensis / Sweltsa kibunensis | KNGDB | | 0 | | 8 | eDNA | | 0 | | 0 | | | 0 | | 0 | |
| Trichoptera | Anisocentropus kawamurai | KNGDB | | 0 | | 0 | | | 0 | | 0 | | | 0 | | 0 | |
| Trichoptera | Nyctiophylax kisoensis | KNGDB | | 0 | | 0 | | | 0 | | 0 | | | 127 | eDNA | 204 | eDNA |
| Trichoptera | Nothopsyche yamagataensis | GenBank | | 0 | | 0 | | | 0 | | 0 | | | 0 | | 0 | |
| Trichoptera | Eobrachycentrus vernalis | KNGDB | | 0 | | 0 | | | 0 | | 0 | | | 0 | | 0 | |
| Trichoptera | Micrasema hanasense | KNGDB | 8 | 0 | Both | 22 | Both | | 0 | | 0 | | 1 | 0 | Capt | 71 | Both |
| Trichoptera | Micrasema uenoi | KNGDB | | 0 | | 0 | | | 0 | | 0 | | | 0 | | 0 | |

| | | | | | | | | | | | | | | | | | | | |
|---|---|---|---|---|---|---|---|---|---|---|---|---|---|---|---|---|---|---|---|
| Trichoptera | Lepidostoma japonicum | KNGDB | | | | 307 | eDNA | 503 | eDNA | 0 | 0 | | | | | 2971 | eDNA | 5879 | eDNA |
| Trichoptera | Lepidostoma naraense | KNGDB | | | | 0 | | 0 | | 0 | 0 | | | | | 32 | eDNA | 112 | eDNA |
| Trichoptera | Lepidostoma orientale | KNGDB | | | | 0 | | 0 | | 0 | 0 | | | | | 0 | | 0 | |
| Trichoptera | Lepidostoma satoi | KNGDB | | | | 0 | | 0 | | 0 | 0 | | | 8 | | 0 | Capt | 0 | Capt |
| Trichoptera | Dolophilodes angustata | KNGDB | | | | 0 | | 0 | | 0 | 0 | | | | | 0 | | 0 | |
| Trichoptera | Dolophilodes iroensis | KNGDB | | | | 0 | | 0 | | 0 | 0 | | | | | 0 | | 0 | |
| Trichoptera | Dolophilodes japonica | KNGDB | | | | 0 | | 0 | | 0 | 0 | | | | | 350 | eDNA | 828 | eDNA |
| Trichoptera | Kisaura minakawai | KNGDB | | | | 8 | eDNA | 14 | eDNA | 0 | 17 | eDNA | | | | 16 | eDNA | 94 | eDNA |
| Trichoptera | Kisaura tsudai | KNGDB | | | | 0 | | 0 | | 0 | 0 | | | | | 0 | | 38 | eDNA |
| Trichoptera | Apsilochorema sutshanum | KNGDB | 6 | 1 | 1 | 57 | Both | 59 | Both | 0 | 10 | eDNA | | | | 240 | eDNA | 325 | eDNA |
| Trichoptera | Psychomyia acutipennis | KNGDB | | | | 0 | | 0 | | 0 | 0 | | | | | 0 | | 0 | |
| Trichoptera | Neophylax japonicus | KNGDB | | | | 0 | | 0 | | 0 | 0 | | 3 | | 10 | 149 | Both | 244 | Both |
| Trichoptera | Uenoa tokunagai | KNGDB | | | | 0 | | 0 | | 0 | 0 | | | | | 0 | | 0 | |

| | | | | | | | | | | | | | | | | | | | | | | | | | | |
|---|---|---|---|---|---|---|---|---|---|---|---|---|---|---|---|---|---|---|---|---|---|---|---|---|---|---|
| Trichoptera | Gumaga orientalis | KNGDB | | | | | 0 | | 0 | | 1 | | | | 0 | Capt | 0 | Capt | | | | | 0 | | 0 | |
| Trichoptera | Apatania aberrans | KNGDB | | | | | 0 | | 56 | eDNA | | | | | 0 | | 0 | | | | | | 442 | eDNA | 525 | eDNA |
| Trichoptera | Cheumatopsyche brevilineata | KNGDB | | | | | 0 | | 0 | | 2 | | | | 955 | Both | 992 | Both | | | | | 0 | | 0 | |
| Trichoptera | Cheumatopsyche infascia | KNGDB | | | | | 0 | | 0 | | | | 4 | | 327 | Both | 574 | Both | | | | | 0 | | 0 | |
| Trichoptera | Diplectrona kibuneana | KNGDB | | | | | 0 | | 0 | | | | | | 0 | | 0 | | | | | | 0 | | 0 | |
| Trichoptera | Hydropsyche albicephala | KNGDB | | | | 1 | 101 | Both | 151 | Both | | | | | 0 | | 0 | | | 1 | 2 | | 145 | Both | 189 | Both |
| Trichoptera | Hydropsyche selysi / Hydropsyche gifuana | KNGDB | | | | | 0 | | 37 | eDNA | | | | | 43 | eDNA | 76 | eDNA | | | | | 0 | | 0 | |
| Trichoptera | Hydropsyche orientalis | KNGDB | 20 | 34 | 5 | 7 | 1097 | Both | 1092 | Both | 29 | 8 | 3 | 2 | 1115 | Both | 2135 | Both | 5 | 1 | 1 | 3 | 813 | Both | 836 | Both |
| Trichoptera | Hydropsyche setensis | KNGDB | | | | | 9 | eDNA | 20 | eDNA | 5 | 7 | 2 | 2 | 3448 | Both | 5823 | Both | | | | | 219 | eDNA | 358 | eDNA |
| Trichoptera | Eubasilissa regina | KNGDB | | | | | 0 | | 0 | | | | | | 0 | | 0 | | | | | | 0 | | 22 | eDNA |
| Trichoptera | Himalopsyche japonica | KNGDB | | | | | 0 | | 10 | eDNA | | | | | 0 | | 0 | | | | | | 0 | | 0 | |
| Trichoptera | Rhyacophila | KNGDB | | | | | 0 | | 0 | | 1 | | | | 31 | Both | 36 | Both | | | | | 0 | | 0 | |

| | | | | | | | | | | | | | | | | | | | | | | | | | | |
|---|---|---|---|---|---|---|---|---|---|---|---|---|---|---|---|---|---|---|---|---|---|---|---|---|---|---|
| | brevicephala | | | | | | | | | | | | | | | | | | | | | | | | | |
| Trichoptera | Rhyacophila clemens | KNGDB | | | | | 0 | | 0 | | | | | | 0 | | 0 | | | | | | 0 | | 0 | |
| Trichoptera | Rhyacophila diffidens | KNGDB | | | | | 0 | | 0 | | | | | | 0 | | 0 | | | | | | 0 | | 0 | |
| Trichoptera | Rhyacophila kawamurae | KNGDB | | | | | 38 | eDNA | 39 | eDNA | | | | | 0 | | 0 | | | | | | 11 | eDNA | 82 | eDNA |
| Trichoptera | Rhyacophila kisoensis | KNGDB | | | | | 0 | | 0 | | | | | | 0 | | 0 | | | | | | 0 | | 30 | eDNA |
| Trichoptera | Rhyacophila towadensis / Rhyacophila lezeyi | KNGDB | | | | | 0 | | 0 | | | | | | 0 | | 0 | | | | | | 0 | | 0 | |
| Trichoptera | Rhyacophila nakagawai | KNGDB | | | | | 0 | | 9 | eDNA | | | | | 0 | | 0 | | | | | | 0 | | 55 | eDNA |
| Trichoptera | Rhyacophila nipponica / Rhyacophila nigrocephala | KNGDB | 6 | 2 | 1 | | 20 | Both | 13 | Both | | | | | 64 | eDNA | 44 | eDNA | 7 | 2 | 4 | 4 | 300 | Both | 212 | Both |
| Trichoptera | Rhyacophila shikotsuensis | KNGDB | | | | | 0 | | 0 | | | | | | 0 | | 0 | | 1 | | | | 0 | Capt | 0 | Capt |
| Trichoptera | Rhyacophila yamanakensis | KNGDB | | | | | 0 | | 0 | | 1 | 2 | 1 | | 155 | Both | 307 | Both | | | | | 0 | | 0 | |

| | | | | | | | | | | | | | | | | | | | | | | | | | | |
|---|---|---|---|---|---|---|---|---|---|---|---|---|---|---|---|---|---|---|---|---|---|---|---|---|---|---|
| Trichoptera | Goera japonica | KNGDB | 2 | | | | 85 | Both | 188 | Both | 2 | | | | 0 | Capt | 0 | Capt | | | | | 116 | eDNA | 357 | eDNA |
| Trichoptera | Stenopsyche marmorata | KNGDB | 3 | 3 | 2 | 1 | 96 | Both | 295 | Both | 95 | 92 | 30 | 17 | 407 | Both | 54 | Both | 1 | | 1 | | 10 | Both | 62 | Both |
| Trichoptera | Mystacides azurea | KNGDB | | | | | 0 | | 0 | | | | | | 0 | | 0 | | | | | | 0 | | 0 | |
| Trichoptera | Hydroptila phenianica | KNGDB | | | | | 0 | | 0 | | | | | | 0 | | 0 | | | | | | 0 | | 0 | |
| Trichoptera | Oxyethira ecornuta | GenBank | | | | | 0 | | 0 | | | | | | 0 | | 0 | | | | | | 0 | | 0 | |
| Trichoptera | Phryganopsyche latipennis | GenBank | | | | | 0 | | 0 | | | | | | 0 | | 0 | | | | | | 0 | | 0 | |
| Trichoptera | Glossosoma ussuricum | KNGDB | | | | | 2114 | eDNA | 3903 | eDNA | | | | | 63 | eDNA | 195 | eDNA | | | | | 3193 | eDNA | 4932 | eDNA |
| Trichoptera | Glossosoma altaicum | KNGDB | | | | | 0 | | 0 | | | | | | 41 | eDNA | 0 | | | | | | 0 | | 0 | |

Database, referenced database used for final species identification; KNGDB, the DNA database for insects created by Kanagawa Prefecture. Capture, the number of insects collected by capture survey; qual, qualitative sampling; quan, quantitative sampling; eDNA, the number of DNA reads detected by eDNA analysis.

**Table S4 Similar sequences to adapters for the forward primer on the genome**

| Species | Accession No. | Scaffold name | Start | End | Query cover | Per ident | Alignment length | E-value |
|---|---|---|---|---|---|---|---|---|
| Trichoptera | | | | | | | | |
| *Limnephilus marmoratus* | GCA_917880875.1 | - | - | - | - | - | - | - |
| *Stenopsyche tienmushanensis* | GCA_008973525.1 | WACJ01000279.1 | 140495 | 140513 | 53 | 94.737 | 19 | 6.8 |
| | | WACJ01000108.1 | 191199 | 191213 | 42 | 100 | 15 | 6.8 |
| | | WACJ01000232.1 | 528381 | 528367 | 42 | 100 | 15 | 6.8 |
| | | WACJ01000159.1 | 686654 | 686640 | 42 | 100 | 15 | 6.8 |
| *Glyphotaelius pellucidus* | GCA_936435175.1 | OW386558.1 | 17587288 | 17587273 | 44 | 100 | 16 | 4 |
| | | OW386539.1 | 9729779 | 9729764 | 44 | 100 | 16 | 4 |
| | | OW386537.1 | 49236551 | 49236532 | 56 | 95 | 20 | 4 |
| Ephemeoroptera | | | | | | | | |
| *Cloeon dipterum* | GCA_949628265.1 | OX451261.1 | 28780222 | 28780236 | 42 | 100 | 15 | 3 |
| *Baetis rhodani* | GCA_001676355.1 | LVVX01110926.1 | 202 | 188 | 42 | 100 | 15 | 2.7 |
| *Ephemera danica* | GCA_000507165.2 | KZ497611.1 | 676868 | 676883 | 44 | 100 | 16 | 1.8 |
| | | AYNC02009728.1 | 22205 | 22219 | 42 | 100 | 15 | 7.2 |
| | | AYNC02007810.1 | 287770 | 287784 | 42 | 100 | 15 | 7.2 |
| | | KZ497891.1 | 296826 | 296812 | 42 | 100 | 15 | 7.2 |
| | | KZ497596.1 | 492606 | 492592 | 42 | 100 | 15 | 7.2 |
| Plecoptera | | | | | | | | |

| | | | | | | | | |
|---|---|---|---|---|---|---|---|---|
| *Protonemura montana* | GCA_947568835.1 | OX387606.1 | 25448234 | 25448251 | 50 | 100 | 18 | 0.067 |
| | | OX387605.1 | 19024366 | 19024382 | 53 | 100 | 17 | 0.26 |
| | | OX387605.1 | 22940395 | 22940410 | 53 | 100 | 16 | 1 |
| | | OX387612.1 | 7983503 | 7983517 | 42 | 100 | 15 | 4.1 |
| | | OX387602.1 | 23511019 | 23511033 | 58 | 100 | 15 | 4.1 |
| | | OX387602.1 | 15336022 | 15336008 | 58 | 100 | 15 | 4.1 |
| | | OX387609.1 | 5703057 | 5703043 | 42 | 100 | 15 | 4.1 |
| | | OX387603.1 | 21804929 | 21804915 | 42 | 100 | 15 | 4.1 |
| *Nemoura dubitans* | GCA_921293005.1 | OV121073.1 | 16959438 | 16959452 | 42 | 100 | 15 | 5.1 |
| | | OV121075.1 | 47690372 | 47690358 | 42 | 100 | 15 | 5.1 |
| *Sweltsa coloradensis* | GCA_024449915.1 | JAMZAV010000080.1 | 1465215 | 1465231 | 47 | 100 | 17 | 1.3 |
| | | JAMZAV010001063.1 | 410753 | 410738 | 44 | 100 | 16 | 5.3 |
| | | JAMZAV010000257.1 | 1759564 | 1759549 | 44 | 100 | 16 | 5.3 |
| *Isoperla grammatica* | GCA_945910005.1 | OX246738.1 | 17931710 | 17931726 | 47 | 100 | 17 | 0.85 |

**Table S5 Similar sequences to adapters for the reverse primer on the genome**

| Species | Accession No. | Scaffold name | Start | End | Query cover | Per ident | Alignment length | E-value |
|---|---|---|---|---|---|---|---|---|
| Trichoptera | | | | | | | | |
| *Limnephilus marmoratus* | GCA_917880875.1 | - | - | - | - | - | - | - |
| *Stenopsyche tienmushanensis* | GCA_008973525.1 | WACJ01000071.1 | 440009 | 440024 | 43 | 100 | 16 | 1.8 |
| | | WACJ01000112.1 | 196398 | 196383 | 43 | 100 | 16 | 1.8 |
| | | WACJ01000109.1 | 596167 | 596153 | 41 | 100 | 15 | 7.2 |
| | | WACJ01000006.1 | 404475 | 404461 | 41 | 100 | 15 | 7.2 |
| *Glyphotaelius pellucidus* | GCA_936435175.1 | OW386544.1 | 21070171 | 21070156 | 43 | 100 | 16 | 4.2 |
| Ephemeoroptera | | | | | | | | |
| *Cloeon dipterum* | GCA_949628265.1 | OX451259.1 | 5700537 | 5700556 | 65 | 95 | 20 | 0.8 |
| | | OX451259.1 | 25144032 | 25144046 | 65 | 100 | 15 | 3.2 |
| | | OX451261.1 | 23498271 | 23498256 | 43 | 100 | 16 | 0.8 |
| | | OX451263.1 | 8442875 | 8442889 | 57 | 100 | 15 | 3.2 |
| | | OX451263.1 | 15911694 | 15911712 | 57 | 94.7 | 19 | 3.2 |
| | | OX451260.1 | 24932904 | 24932918 | 41 | 100 | 15 | 3.2 |
| *Baetis rhodani* | GCA_001676355.1 | LVVX01381374.1 | 4256 | 4270 | 41 | 100 | 15 | 2.8 |
| | | LVVX01243396.1 | 45 | 63 | 51 | 94.7 | 19 | 2.8 |
| *Ephemera danica* | GCA_000507165.2 | KZ498009.1 | 524330 | 524351 | 59 | 100 | 22 | 5.01E-04 |
| | | KZ497760.1 | 89740 | 89720 | 57 | 100 | 21 | 0.002 |

| | | | | | | | | |
|---|---|---|---|---|---|---|---|---|
| | | KZ498429.1 | 15571 | 15544 | 76 | 92.9 | 28 | 0.008 |
| | | KZ497819.1 | 90729 | 90759 | 84 | 90.3 | 31 | 0.031 |
| | | KZ497839.1 | 105024 | 105006 | 51 | 100 | 19 | 0.031 |
| | | KZ498729.1 | 8096 | 8112 | 46 | 100 | 17 | 0.48 |
| | | KZ501762.1 | 1271065 | 1271080 | 43 | 100 | 16 | 1.9 |
| | | KZ497647.1 | 438125 | 438140 | 43 | 100 | 16 | 1.9 |
| | | KZ497638.1 | 48142 | 48165 | 65 | 91.7 | 24 | 1.9 |
| | | KZ498120.1 | 81414 | 81399 | 43 | 100 | 16 | 1.9 |
| | | KZ497755.1 | 360599 | 360584 | 43 | 100 | 16 | 1.9 |
| Plecoptera | | | | | | | | |
| *Protonemura montana* | GCA_947568835.1 | OX387612.1 | 6574705 | 6574690 | 57 | 100 | 16 | 1.1 |
| | | OX387612.1 | 11444054 | 11444068 | 57 | 100 | 15 | 4.3 |
| | | OX387605.1 | 15941354 | 15941339 | 43 | 100 | 16 | 1.1 |
| *Nemoura dubitans* | GCA_921293005.1 | OV121075.1 | 12703352 | 12703367 | 49 | 100 | 16 | 1.3 |
| | | OV121075.1 | 15907258 | 15907272 | 49 | 100 | 15 | 5.1 |
| | | OV121073.1 | 62063870 | 62063855 | 57 | 100 | 16 | 1.3 |
| | | OV121073.1 | 36352337 | 36352323 | 57 | 100 | 15 | 5.1 |
| *Sweltsa coloradensis* | GCA_024449915.1 | JAMZAV010014428.1 | 72 | 91 | 54 | 100 | 20 | 0.023 |
| | | JAMZAV010007108.1 | 19648 | 19663 | 43 | 100 | 16 | 5.6 |
| | | JAMZAV010002591.1 | 63455 | 63470 | 43 | 100 | 16 | 5.6 |
| | | JAMZAV010001060.1 | 46490 | 46505 | 43 | 100 | 16 | 5.6 |
| | | JAMZAV010003266.1 | 136469 | 136454 | 43 | 100 | 16 | 5.6 |

| | | | | | | | | |
|---|---|---|---|---|---|---|---|---|
| | | JAMZAV010000070.1 | 257601 | 257586 | 43 | 100 | 16 | 5.6 |
| *Isoperla grammatica* | GCA_945910005.1 | OX246741.1 | 15631948 | 15631963 | 43 | 100 | 16 | 3.5 |
| | | OX246738.1 | 12667483 | 12667468 | 43 | 100 | 16 | 3.5 |

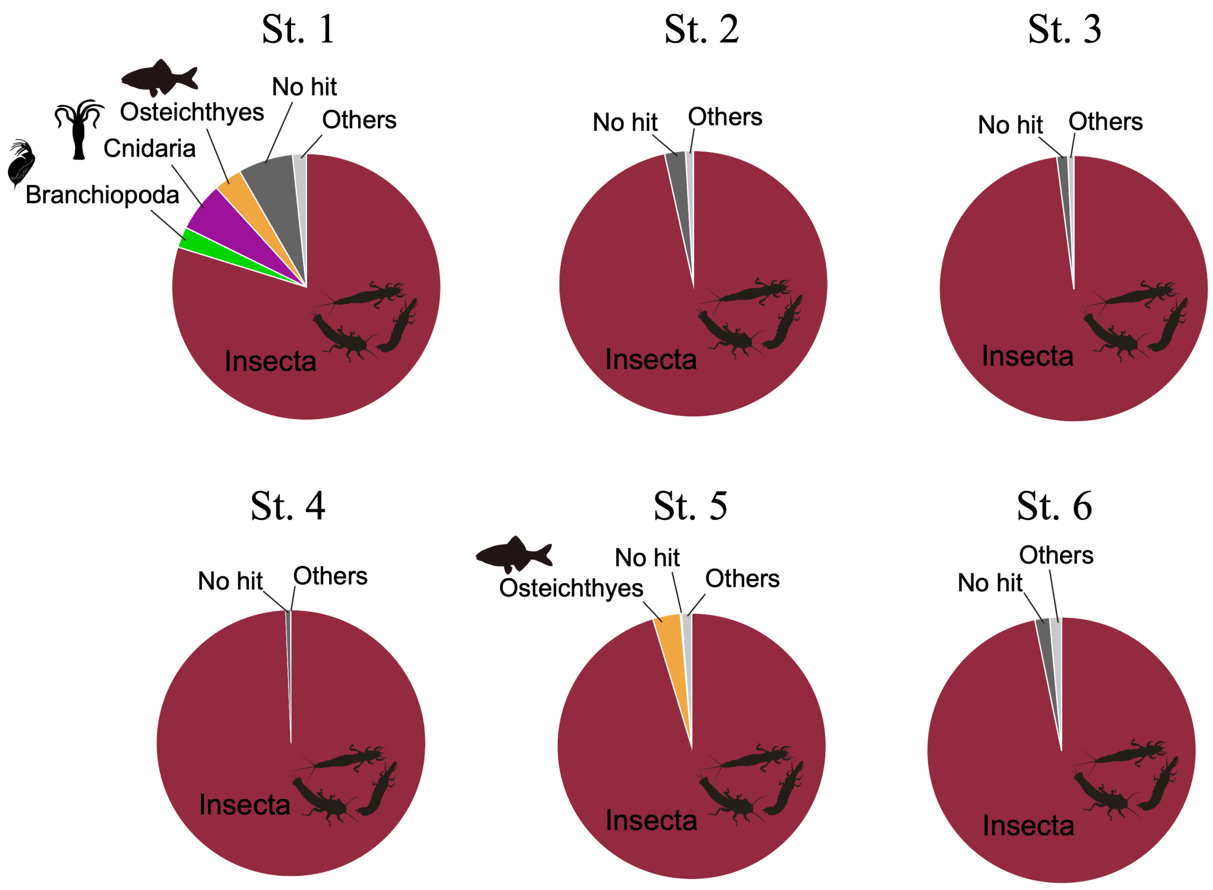

Figure S1. Pie charts showing the percentage of reads detected from eDNA analyses of major animal taxa at each site. No hit, eDNA reads that did not hit the database; Others, detecting eDNA read of other groups.

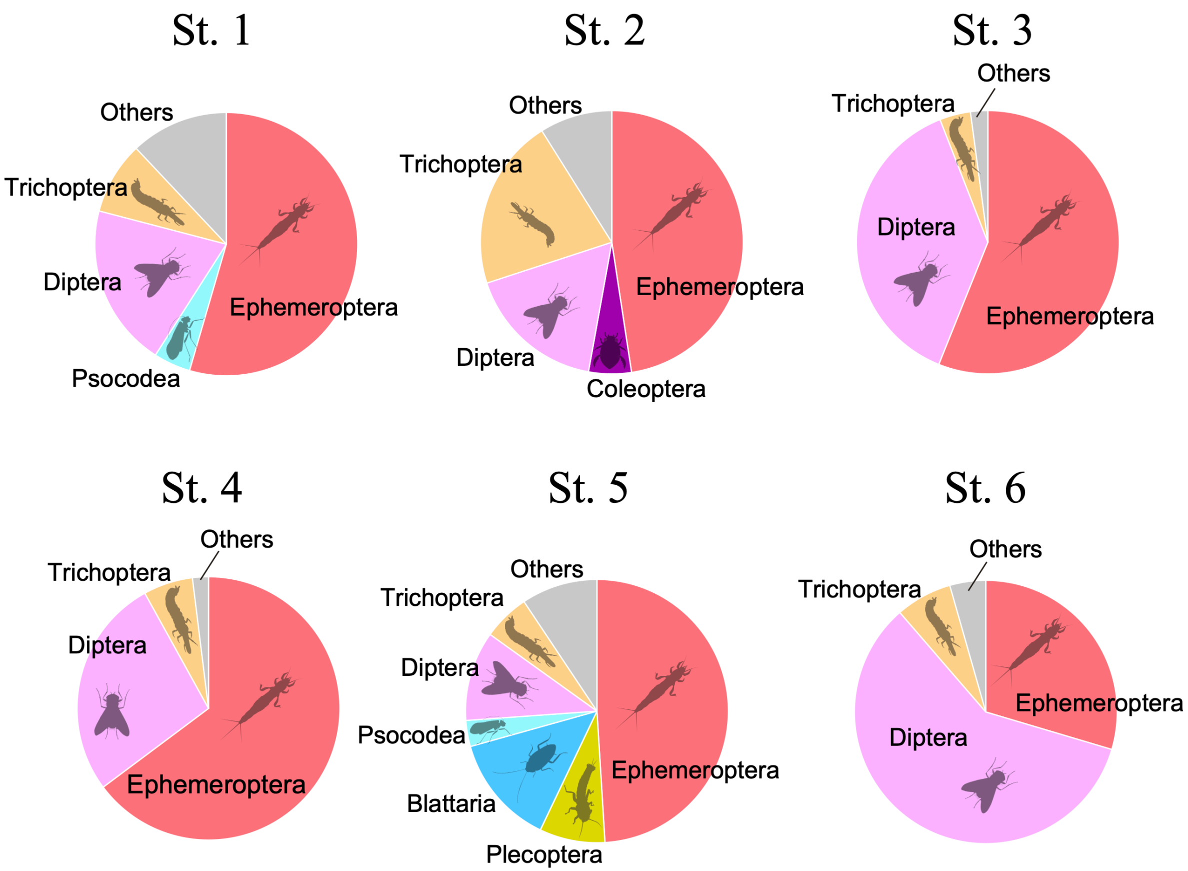

Figure S2. Pie charts showing the percentage of reads detected from eDNA analyses at order level within Insecta.

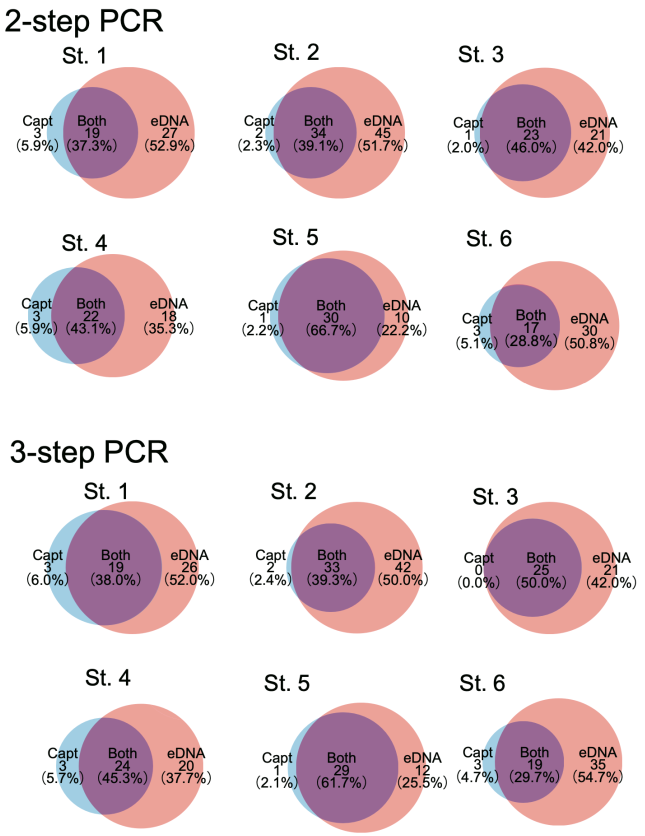

Figure S3. Venn diagrams showing the comparison of the number of species detected by eDNA analyses and actual capture surveys, and the comparison of two library prep patterns.

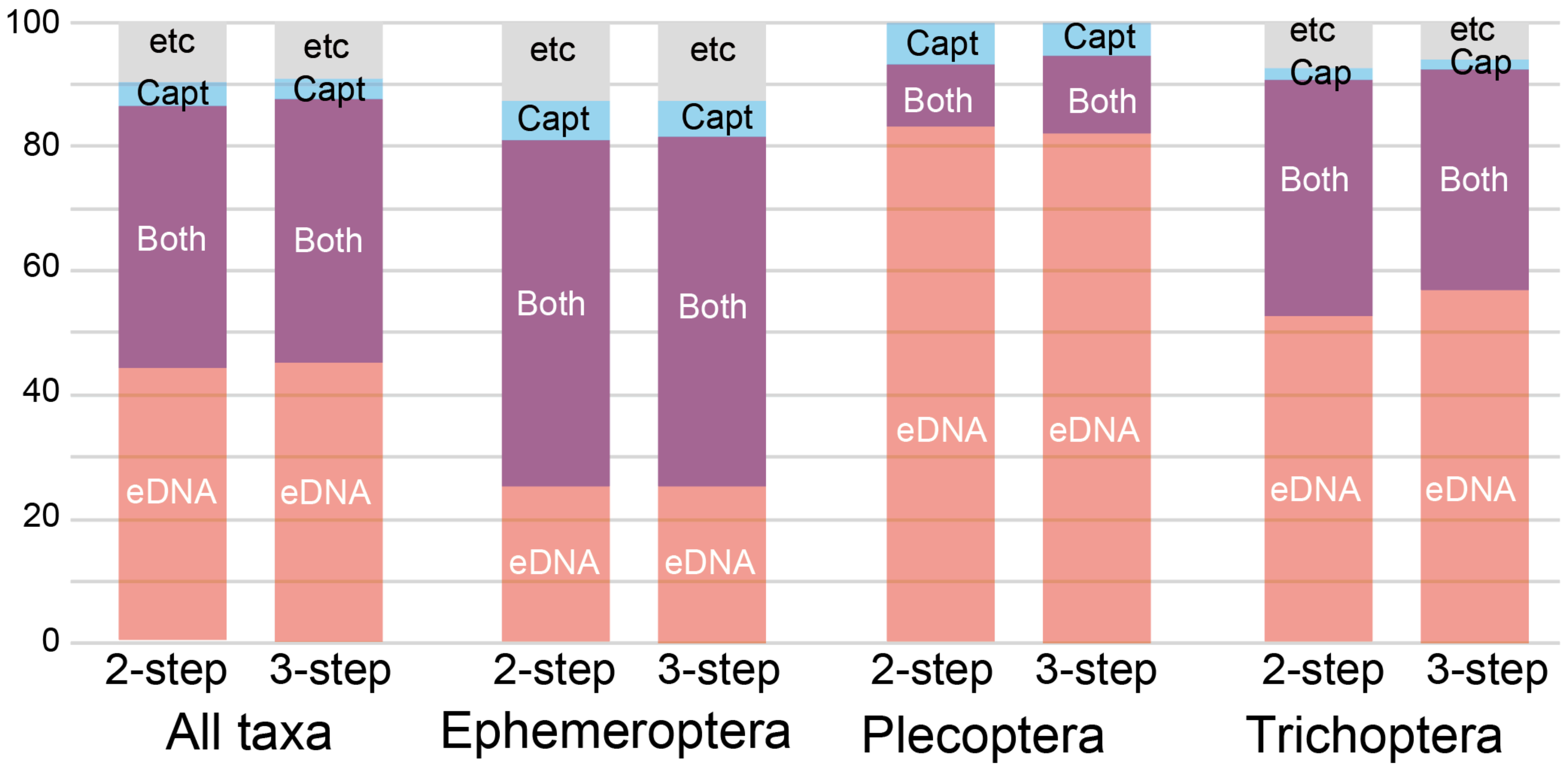

Figure S4. Comparison of the percentage of species detected by eDNA analyses and actual capture surveys within each major order, Ephemeroptera, Plecoptera, and Trichoptera, and comparison of two library prep patterns, the 2-step PCR and 3-step PCR methods. eDNA, the percentage of species detected by eDNA analyses; Capt, the percentage of species detected by actual capture surveys; Both, the percentage of species detected by both eDNA analyses and actual capture surveys; etc, eDNA reads that did not hit the database or treated as non-data. The number of species detected is shown in Table S3.
